# Biomolecular solid-state NMR spectroscopy at highest field: the gain in resolution at 1200 MHz

**DOI:** 10.1101/2021.03.31.437892

**Authors:** Morgane Callon, Alexander A. Malär, Sara Pfister, Václav Rímal, Marco E. Weber, Thomas Wiegand, Johannes Zehnder, Matías Chávez, Rajdeep Deb, Riccardo Cadalbert, Alexander Däpp, Marie-Laure Fogeron, Andreas Hunkeler, Lauriane Lecoq, Anahit Torosyan, Dawid Zyla, Rudolf Glockshuber, Stefanie Jonas, Michael Nassal, Matthias Ernst, Anja Böckmann, Beat H. Meier

## Abstract

Progress in NMR in general and in biomolecular applications in particular is driven by increasing magnetic-field strengths leading to improved resolution and sensitivity of the NMR spectra. Recently, persistent superconducting magnets at a magnetic field strength (magnetic induction) of 28.2 T corresponding to 1200 MHz proton resonance frequency became commercially available. We present here a collection of high-field NMR spectra of a variety of proteins, including molecular machines, membrane proteins and viral capsids and others. We show this large panel in order to provide an overview over a range of representative systems under study, rather than a single best performing model system. We discuss both carbon-13 and proton-detected experiments, and show that in ^13^C spectra substantially higher numbers of peaks can be resolved compared to 850 MHz while for ^1^H spectra the most impressive increase in resolution is observed for aliphatic side-chain resonances.

## Introduction

New technologies have often stood at the beginning of new spectroscopic techniques and NMR is a particularly good example: Microcomputers have enabled Fourier spectroscopy (Ernst and Anderson 1965) and multidimensional NMR (Aue, Bartholdi, and Ernst 1976), high and stable magnetic fields generated by persistent superconducting magnets have been instrumental for the first protein structure determinations (Wüthrich 2003; Williamson, Havel, and Wuthrich 1985) and the structural and dynamic investigation of increasingly larger proteins (Rosenzweig and Kay 2014; Pervushin et al. 1997; Fiaux et al. 2002). Reliable magic-angle sample spinning probes together with high magnetic fields have enabled biomolecular solid-state NMR spectroscopy (McDermott et al. 2000). The first solid-state NMR protein-structure determination used a magnetic-field strength of 17.6 T (proton resonance frequency 750 MHz) (Castellani et al. 2002), and the first prion fibril structure was determined at 850 MHz (Wasmer et al. 2008). A next achievement with important impact was the development of fast magic-angle spinning (MAS) probes, in excess of 100 kHz rotation frequency, enabling proton detection and a reduction of the required sample amount by roughly two orders of magnitude (Agarwal et al. 2014; Andreas et al. 2015; Barbet-Massin et al. 2014; Schledorn et al. 2020; Penzel et al. 2019; Lecoq et al. 2019).

Since 1000 MHz proton Larmor frequency is the present limit of what could be achieved with low-temperature superconducting (LTS) wire (such as Nb3Sn and NbTi), persistent magnetic fields exceeding 1000 MHz required solenoid coils made out of high-temperature superconducting (HTS) wire (e.g. REBCO) (Maeda and Yanagisawa 2019). Thus, after the highest LTS magnet (1 GHz), it has taken more than five years to develop this new technology and achieve higher fields. Today, persistent hybrid superconducting magnets combining both, LTS and HTS, have been developed by Bruker Switzerland AG generating magnetic-field strengths up to 28.2 T corresponding to 1200 MHz proton Larmor frequency.

What improvement in resolution and sensitivity do we expect by an increase in magnetic field from 850 to 1200 MHz? Assuming that the NMR linewidths are dominated by scalar couplings or residual dipolar couplings under MAS, they should be field-independent when expressed in frequency units (Hz). Then, resolution in NMR spectra benefits when going from 850 to 1200 MHz through an increase in chemical-shift dispersion (in Hz) by a factor of nearly 1.5 (the ratio of the two magnetic fields). On the ppm scale, the linewidth decreases linearly with increasing *B*_0_ by the same factor of around 1.5 (see Figure S1 for an illustration). With respect to sensitivity, the theoretical gain in signal-to-noise ratio (SNR) is given by 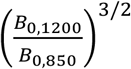 (Abragam 1961), which corresponds to a factor of 1.7 in the integral of the peaks. These considerations apply both to ^13^C- and ^1^H-detected experiments.

The above values are valid for “perfect” samples, which do neither show conformational disorder (resulting in heterogenous line broadening), nor dynamics (resulting in homogenous line broadening). Heterogeneous line broadening scales up linearly with the magnetic field. This contribution to the total linewidth is independent of *B*_0_, and stays constant in ppm. In real samples, both disorder and dynamics can represent important contributions to the linewidths; this is why it is important to illustrate the gain achieved for a broad selection of samples. Besides these sample-dependent effects, several instrumental imperfections can limit the quality of the spectra, including magnetic-field inhomogeneity in space (shims) and in time (field drifts), or imperfect or unstable magic-angle adjustment and radiofrequency field (rf) inhomogeneity. There are a number of intrinsic challenges when going to higher fields: the larger chemical-shift dispersion makes the application of higher power pulses necessary to cover the entire spectrum; at the same time, obtaining high radio-frequency (rf) fields becomes more demanding at higher frequency, in particular for lossy samples with a high salt content.

We herein present first results obtained on a 1200 MHz spectrometer for a set of biomolecular samples that we have already investigated at 850 MHz, and compare sensitivity and resolution in ^1^H- and ^13^C-detected NMR spectra. We avoided the temptation to select one “typical” sample, i.e. the very best performing sample that we have, but rather present a selection of samples that we are currently investigating in the laboratory. We used both, the more classical approach of ^13^C-detected spectroscopy, which is of advantage when large sample quantities (approx. 30 mg) can be prepared, as well as ^1^H-detected solid-state NMR, which has a mass sensitivity about 50 times higher, and relies on the use of sub milligram protein quantities (Lecoq et al. 2019; Agarwal et al. 2014). Both approaches are today central in biomolecular NMR spectroscopy, and show different strengths and limitations. Proton-detected spectra at 1200 MHz are also under investigation in other labs (Nimerovsky et al. 2021).

## Results

In the following, we compare spectra of amyloid fibrils of the fungal prion HET-s(218-289) (Wasmer et al. 2008; van Melckebeke et al. 2010); sediments of the bacterial helicase DnaB (Gardiennet et al. 2012; Wiegand et al. 2019); the bacterial RNA helicase and acetyltransferase TmcA (Ikeuchi, Kitahara, and Suzuki 2008; Chimnaronk et al. 2009); the Rpo4/7 protein complex of two subunits of archeal RNA polymerase II (Torosyan et al. 2019); the filaments of PYRIN domain of mouse ASC (Sborgi et al. 2015; Ravotti et al. 2016); the viral capsids of the Hepatitis B virus (Lecoq et al. 2019) and the African cichlid nackednavirus (Lauber, Seitz at al. 2017); supramolecular protein filaments of type 1 pili (Hahn et al. 2002; Habenstein et al. 2015); and the nonstructural membrane protein 4B (NS4B) of the Hepatitis C virus. In all figures, spectra colored in blue were recorded at 850 MHz, and spectra in red at 1200 MHz.

### ^13^C-detected ^13^C-^13^C correlation spectroscopy

Figure 1a and b compare ^13^C- and ^15^N-detected cross-polarization (CP) spectra of HET-s(218-289) amyloid fibrils (Wasmer et al. 2008; van Melckebeke et al. 2010; Smith et al. 2017) measured in a commercial Bruker 3.2mm triple-resonance probe using the E-free design (Gor’kov et al. 2007) at 850 and 1200 MHz. Out of the 71 residues of HET-s(218-289), 56 are observed in CP spectra (van Melckebeke et al. 2010), the remainder is invisible due to dynamics (Smith 2018; Siemer et al. 2006). A significant sensitivity gain is observed in the ^13^C spectrum at 1200 MHz, although not homogeneous over all resonances, but the aliphatic region is favored. We attribute this inhomogeneity of the sensitivity gain to the offset dependence of the CP step caused by the limited rf-field strength available at the 1200 MHz spectrometer on the ^13^C channel of the probe (the ~48 kHz used are not large compared to the ^13^C spectral width). The aliphatic regions of 2D ^13^C-^13^C Dipolar Assisted Rotational Resonance (DARR) spectra (Takegoshi, Nakamura, and Terao 2001; 2003) of HET-s(218-289) recorded at 850 and 1200 MHz are given in Figure 1c, with expanded regions shown in Figure 1d. It can clearly be seen that the spectra at 1200 MHz show higher resolution. However, one can conclude from the 1D traces (Figure S3) that the DARR transfer is, as expected, somewhat less efficient for constant mixing time at the higher magnetic field. Since the MAS frequency of the 1200 MHz 3.2mm probe is currently limited to 20 kHz, corresponding to ~66 ppm, some rotational-resonance (Raleigh, Levitt, and Griffin 1988; Colombo, Meier, and Ernst 1988) line-broadening effects are present at 1200 MHz between the carbonyl and aliphatic resonances. The linewidth thus can still be improved by spinning faster; a MAS frequency of around 24 kHz would be optimal, and can generally be achieved in 3.2 mm rotors (Böckmann, Ernst, and Meier 2015).

**Figure 1:**
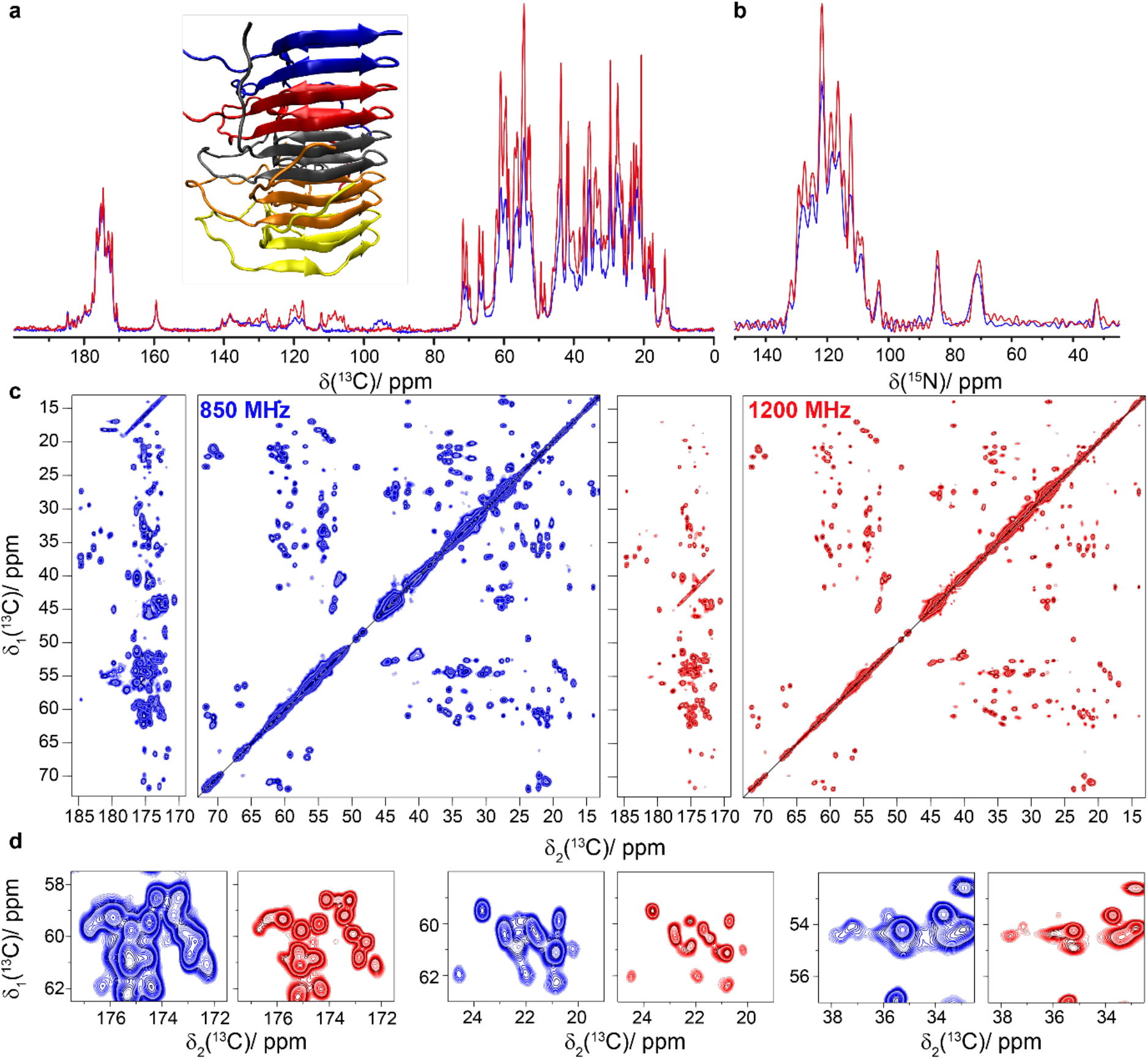
HET-s(218-289) amyloid fibrils. **a)** Structure model (PDB ID: 2RNM)(Wasmer et al. 2008) and 1D ^13^C-detected CP-MAS spectrum, **b)** 1D ^15^N-detected CP-MAS spectrum, **c)** 20 ms DARR spectra and **d)** expanded regions from the spectra in **c**. Spectra colored in blue were recorded at 850 MHz and spectra in red were measured at 1200 MHz. CP was matched at 75 and 48 kHz for ^1^H and ^13^C at 1200 MHz and 60 and 43 kHz at 850 MHz. Experimental parameters are listed in Table S1. The two spectra were normalized using isolated well-resolved peaks and the contour levels are the same for the two spectra.

As a second system, we show adhesive type 1 pili from *E.coli*, which assemble *in vitro* to form long supramolecular protein filaments (Hahn et al. 2002; Habenstein et al. 2015). Each monomer consists of 150 amino acids. Using 850 MHz data, the ^13^C resonances have been assigned to 98% of the sequence (Habenstein et al. 2015). A clear improvement in resolution at the higher field is observed in the expanded regions shown in Figure2c.

**Figure 2:**
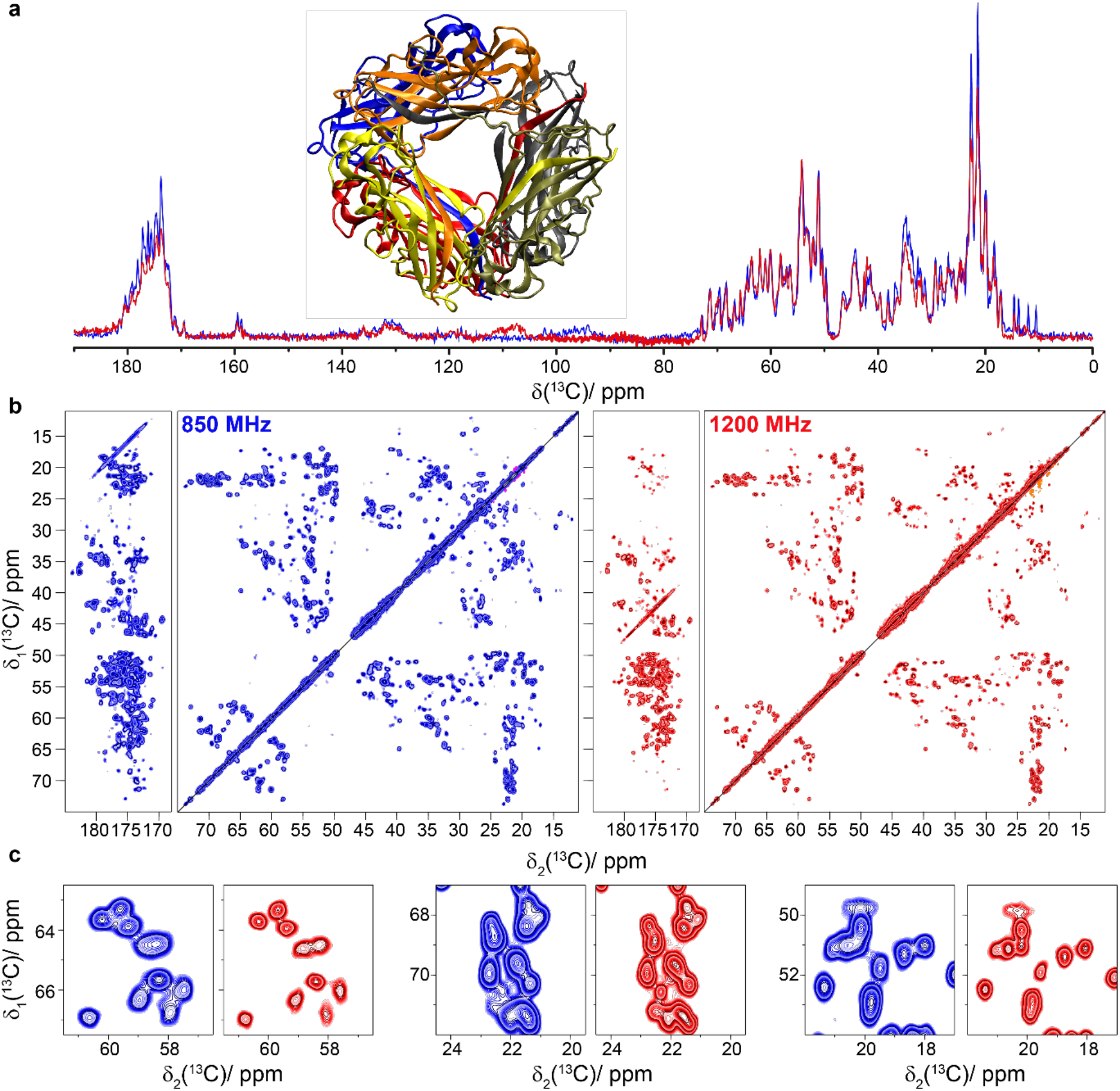
Protein filaments of type 1 pili. **a)** Structural model (PDB ID: 2N7H) (Habenstein et al. 2015) and 1D ^13^C-detected CP-MAS spectrum, **b)** 20 ms DARR spectra and **c)** spectral fingerprints expanded from the spectra in **b**. Spectra colored in blue were recorded at 850 MHz and spectra in red were measured at 1200 MHz. CP was matched at 70 and 44 kHz for ^1^H and ^13^C at 1200 MHz and at 60 and 43 kHz at 850 MHz.

While the type 1 pili and HET-s(218-289) were small enough for assignment and structure determination at 850 MHz (Wasmer et al. 2008; van Melckebeke et al. 2010) (Habenstein et al. 2015), the DnaB helicase from *Helicobacter pylori* (6 x 59 kDa) with 488 residues per monomer poses a big challenge at 850 MHz, already for assignment. Divide-and-conquer approaches (Wiegand, Gardiennet, Cadalbert, et al. 2016; Wiegand, Gardiennet, Ravotti, et al. 2016) or segmental isotope labeling of individual protein domains have thus been applied (Wiegand 2018), however without reaching close-to-complete assignment. Figure 3 shows the NMR spectra collected on DnaB complexed with ADP:AlF_4_^-^ and single-stranded DNA (Wiegand et al. 2019). Figure 3a displays the ^13^C-detected 1D CP-spectra recorded at three different magnetic field strengths: 500 MHz, 850 MHz and 1200 MHz. The efficiency of the CP at 1200 MHz suffers again from offset effects, even more than in Figure 1a, as the higher salt content of the sample (130 mM NaCl) reduces the rf-field strengths that can be safely applied according to the manufacturer. Figure 3b shows the 20 ms DARR spectra recorded at these different magnetic field strengths. The increase in resolution with magnetic field is obvious (Figure 3b,c). We have automatically picked the resonances in the aliphatic region (using CCPNmr (Fogh et al. 2002; Vranken et al. 2005)) of the 2D spectra and find an increase from 203 to 322 peaks between the 850 and 1200 MHz spectra (see Figure S4), highlighting the gain in resolution. This gain will allow us to go further in structural studies of this protein using 3D and 4D spectra to further increase resolution.

**Figure 3:**
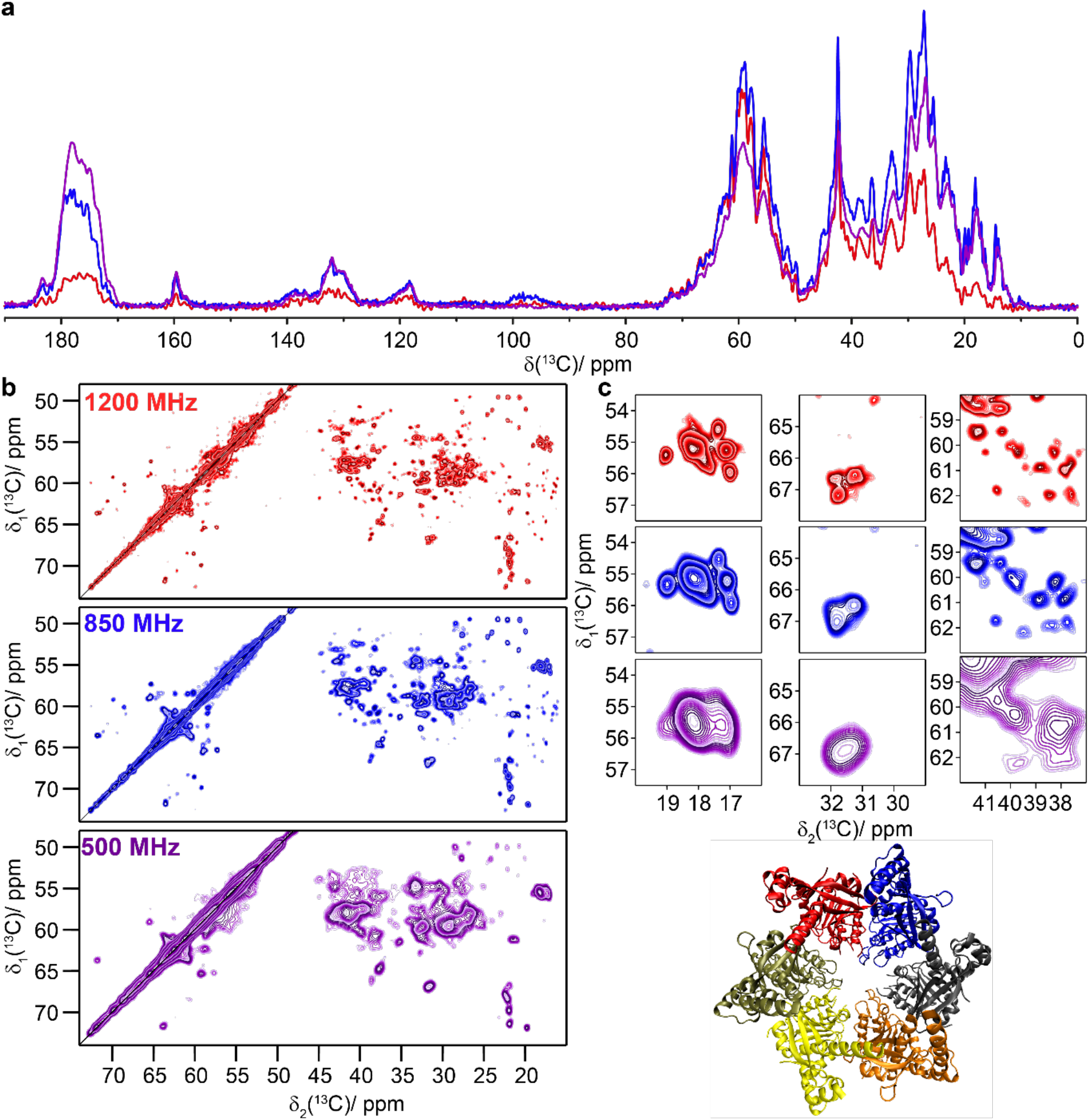
The bacterial DnaB helicase. **a)** 1D ^13^C-detected CP-MAS spectra recorded at 500, 850, and 1200 MHz; **b)** 20 ms DARR spectra recorded at the same magnetic fields as in **a**. **c)** Expanded regions from the spectra in **b**. Spectra colored in purple were measured at 500 MHz, spectra in blue were recorded at 850 MHz and spectra in red were measured at 1200 MHz. CP was matched at 55 and 29 kHz for ^1^H and ^13^C at 1200 MHz and at 60 and 43 kHz at 500 and 850 MHz. The 1D spectra in **a** were scaled to a similar noise level. Structural model with each subunit colored differently (PDB ID: 4ZC0) (Bazin et al. 2015).

As an example for an even larger protein with 671 amino-acid residues, we studied the RNA helicase and acetyltransferase TmcA (Figure 4) (Ikeuchi, Kitahara, and Suzuki 2008; Chimnaronk et al. 2009). 3.2mm E-free probe and experiments at 1200 MHz might open up the possibility of at least partial assignments as suggested by the demonstrated gain in resolution.

**Figure 4:**
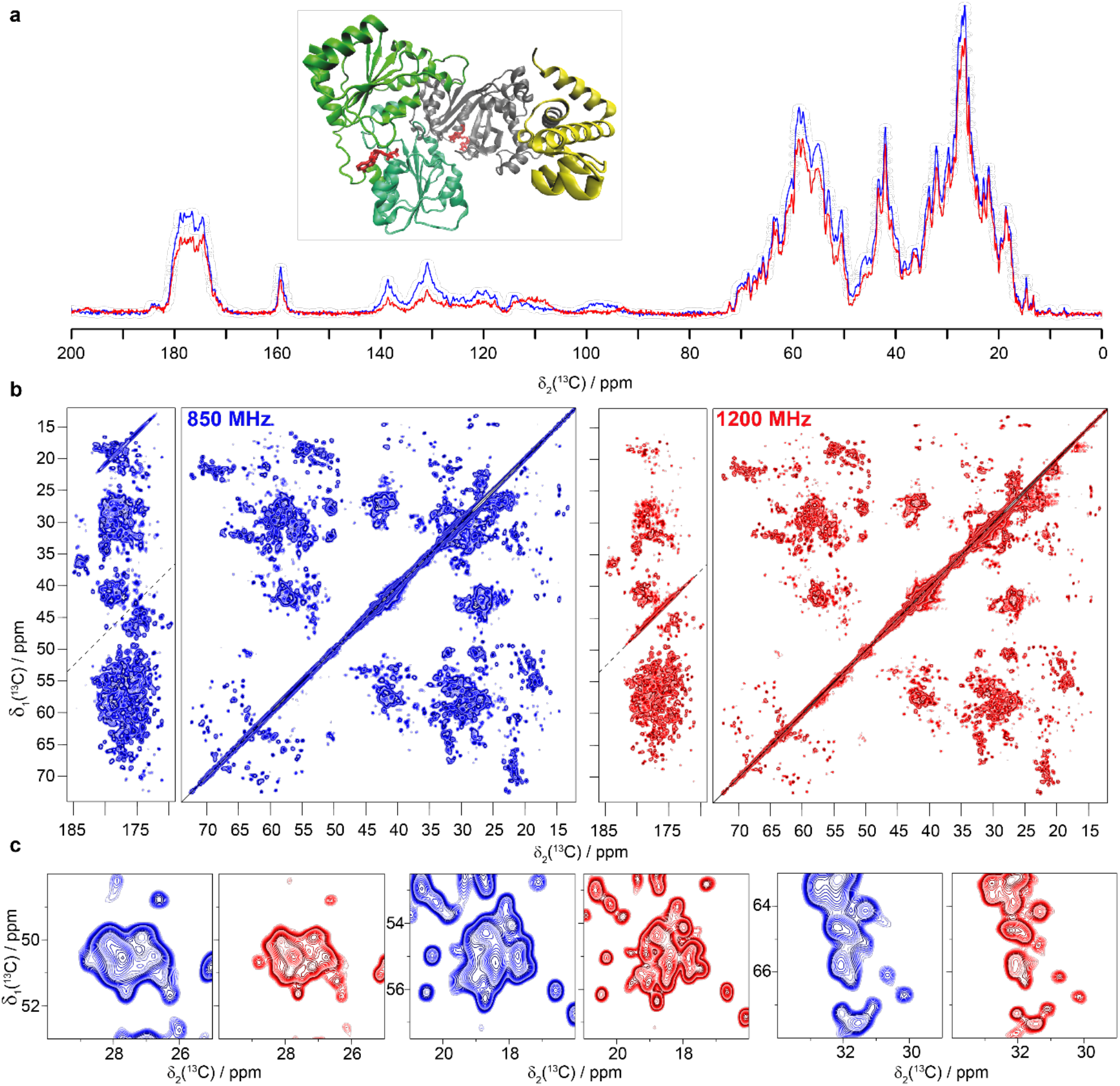
The RNA helicase and acetyltransferase TmcA. **a)** Structural model (PDB ID: 2ZPA) (Chimnaronk et al. 2009) and 1D ^13^C-detected CP-MAS spectrum, **b)** 20 ms DARR spectra and **c)** expanded regions from the spectra in **b**. Spectra colored in blue were recorded at 850 MHz and spectra in red were measured at 1200 MHz. CP was matched at 70 and 44 kHz for ^1^H and ^13^C at 1200 MHz and at 60 and 43 kHz at 850 MHz.

Finally, we measured the CP and DARR spectra of the viral capsid of the African chichlid nackednavirus, a non-enveloped fish virus and member of the Hepatitis B family (Lauber, Seitz et al. 2017). The core protein constituting the capsid consists of 175 amino-acid residues. The corresponding spectra are shown in Figure 5. The T=3 icosahedral capsid is formed by 60 copies of A, B and C subunits that constitute the asymmetric unit. Unlike in the case of the HBV capsid (Lecoq et al. 2018), the signals from the different subunits are not resolved in the nackednavirus capsid spectra. They may, however, contribute to the heterogeneous line broadening that limits the resolution improvement when going to the higher field. As a consequence, the gain in spectral resolution is somewhat less spectacular than for the other samples.

**Figure 5:**
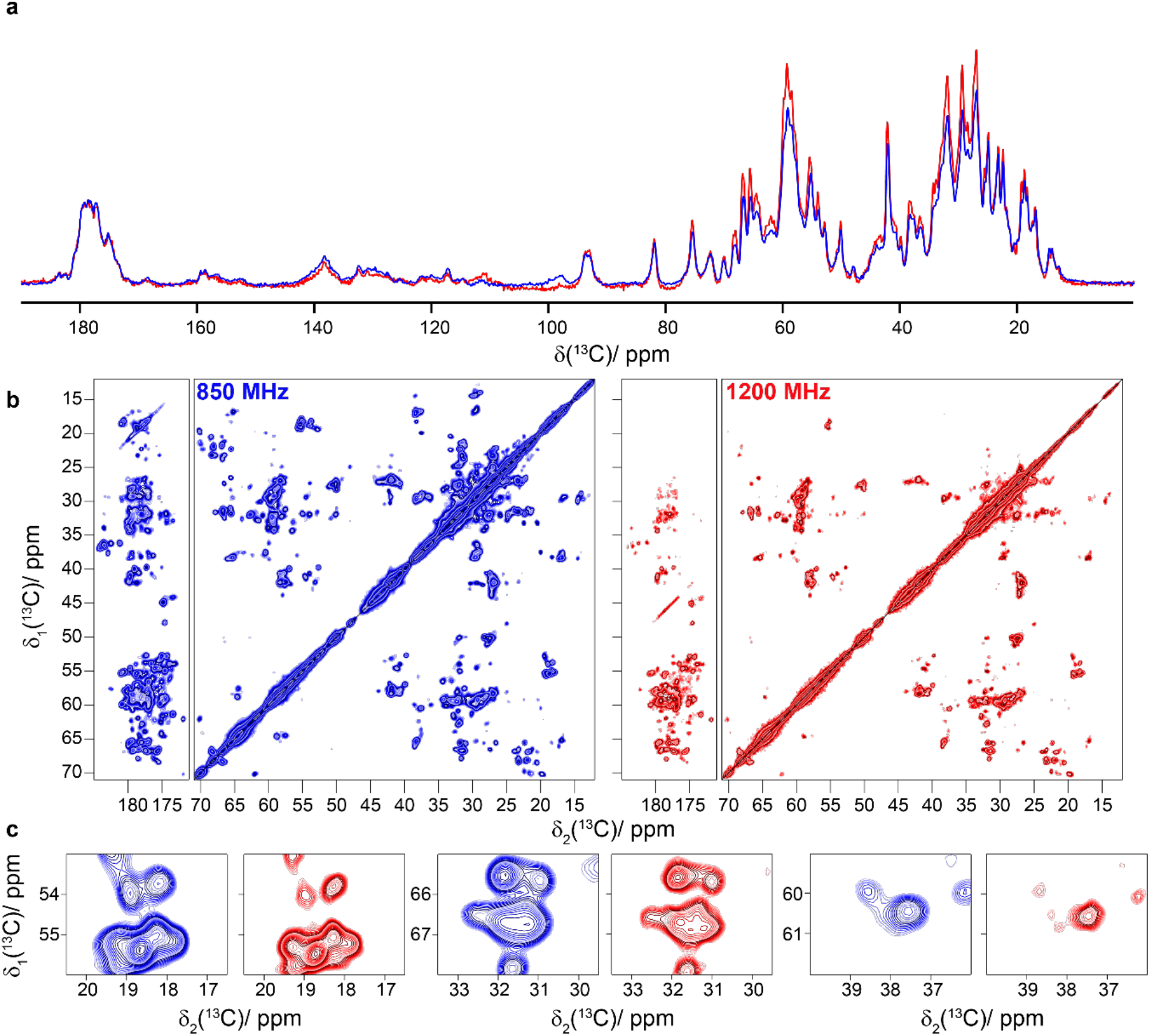
The African cichlid nackednavirus capsid ACNDVc. **a)** 1D ^13^C-detected CP-MAS spectrum, **b)** 20 ms DARR spectra and **c)** expanded regions from the spectra in **b**. Spectra colored in blue were recorded at 850 MHz and spectra in red were measured at 1200 MHz. CP was matched at 75 and 47 kHz for ^1^H and ^13^C at 1200 MHz and at 60 and 41 kHz at 850 MHz.

### ^1^H-detected ^1^H-^15^N correlation spectroscopy

In addition to faster spinning (Penzel et al. 2019; Schledorn et al. 2020), higher fields can equally improve proton resolution(Xue et al. 2020). In samples where the linewidth is dominated by coherent homogeneous interactions, one can expect a linear improvement in resolution. Importantly, beyond this, further narrowing can be induced by the truncation of strong-coupling effects by the increased chemical-shift difference between the coupled protons at higher magnetic fields.

Strong coupling effects are important if the chemical-shift difference between two spins *m* and *n* is smaller than the second-order residual dipolar terms that include *m* and *n* (Malär et al. 2019). In this case, the two resonances are not clearly separated within the dipolar linewidth. This effect is often most relevant for CH_2_ groups which have a strong dipolar coupling, and the chemical shifts of neighboring CH_2_ protons are much closer than for example H_N_ or H*α* protons.

We first investigated the Hepatitis B virus (HBV) nucleocapsid, composed of 240 copies of the core protein (Cp) (Wynne, Crowther, and Leslie 1999; Lecoq et al. 2019). Cp149 is a truncated version containing only the assembly domain. The 2D hNH spectra of ^2^H-^13^C-^15^N labeled and re-protonated Cp149 (dCp149) capsids are shown in Figure 6. We note a small increase in sensitivity in the one-dimensional ^1^H-detected spectrum at 1200 MHz (Figure 6a); more importantly, the increase in resolution at the higher field is very clear. This can be visualized by comparing the 2D hNH spectra and expanded regions shown in Figures 6c and d, respectively, as well as in the one-dimensional trace at δ_1_(^15^N) =118.5 ppm (Figure 6b). Peak picking performed in the amide region of the 2D spectra shows an increase from 110 to 157 peaks when going from 850 MHz to 1200 MHz (see Figure S5). As discussed for the ^13^C spectra of the nackednavirus, the resolution may be limited by unresolved peak splittings due to the presence of four molecules per asymmetric unit (Lecoq et al. 2019).

**Figure 6:**
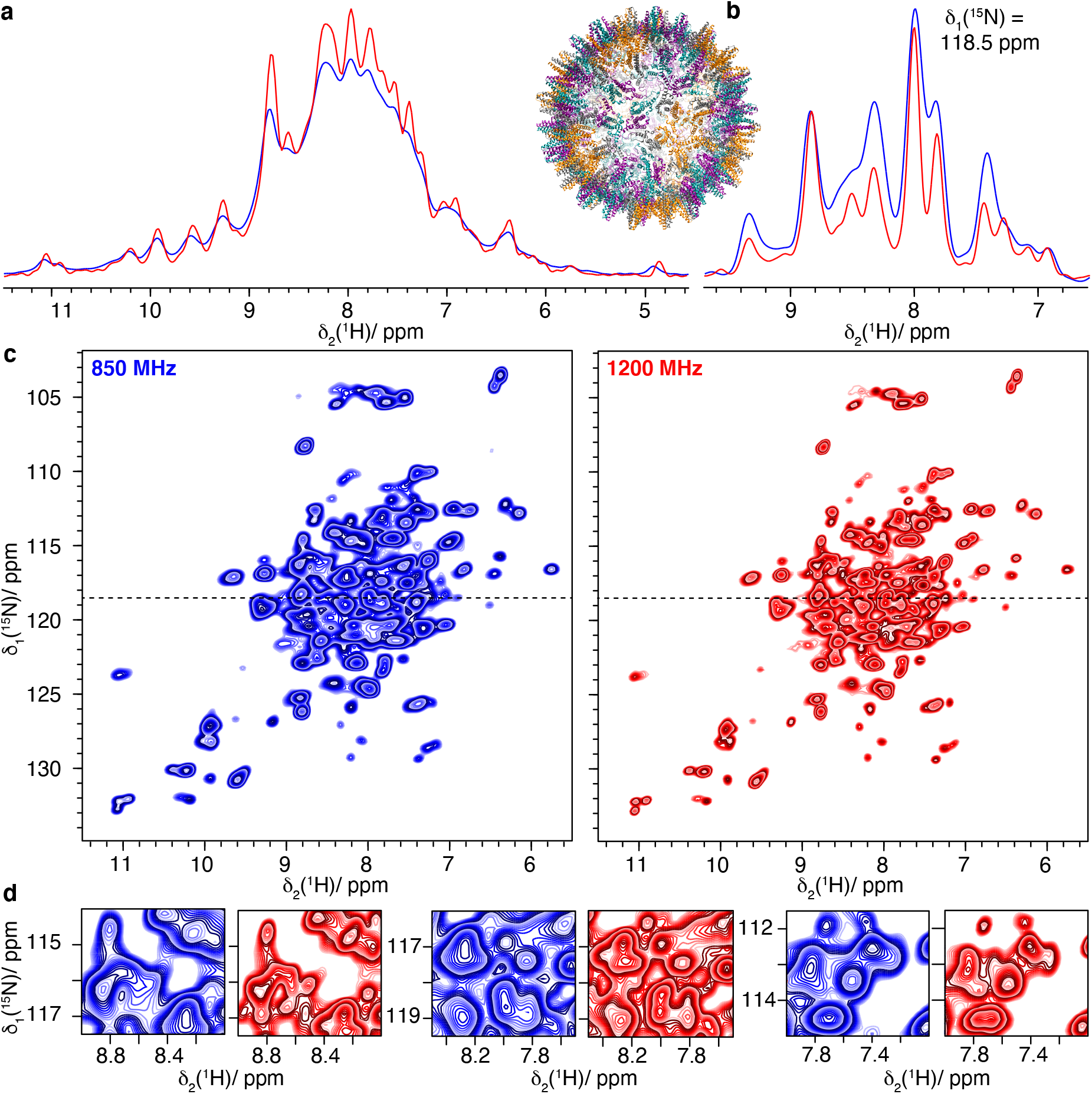
The Hepatitis B Virus Capsid dCp149. **a)** 1D-hnH spectra and structural model (PDB ID:1QGT) (Wynne, Crowther, and Leslie 1999), **b)** one-dimensional trace at δ_1_(^15^N)=118.5 ppm of **c)** 2D hNH spectra and **d)** expanded regions from the spectra in **c**. Spectra colored in blue were recorded at 850 MHz and spectra in red were measured at 1200 MHz.

Investigations of integral membrane proteins in lipids still remain at the very edge of what is possible at current magnetic fields, because of their poor chemical shift dispersion due to their mainly *α*-helical secondary structure. We recently studied the cell-free synthesized hepatitis C virus (HCV) nonstructural membrane protein 4B (NS4B) (Jirasko et al. 2020). NS4B has a sequence length of 261 amino-acid residues, and is an oligomeric *α*-helical integral membrane protein constituted of three subdomains (Gouttenoire et al. 2014). Even though multidimensional spectra could be obtained, sequential assignments could only be achieved for few residues (Jirasko et al. 2020).

Figure 7 shows the 2D hNH spectra of ^2^H-^13^C-^15^N labeled NS4B (dNS4B). A clear gain, both in sensitivity and resolution, is observed at 1200 MHz. This gain will enable further steps in the backbone assignment using 3D approaches, and might allow secondary-structure analysis sufficient for a critical evaluation of existing models.

**Figure 7:**
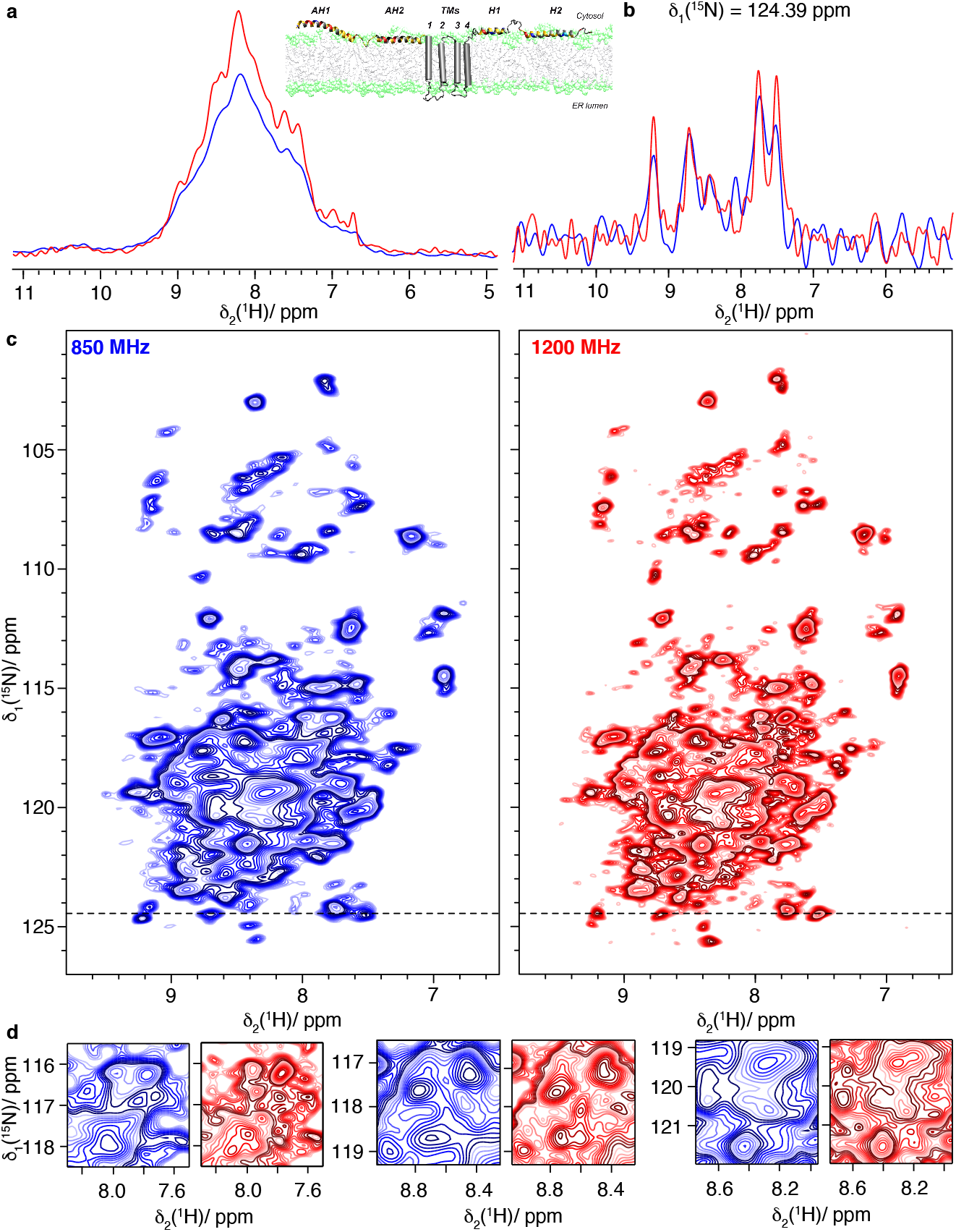
The hepatitis C virus non-structural protein dNS4B. **a)** 1D-hnH spectra and structural model based on (Gouttenoire et al. 2014), **b)** one-dimensional trace at δ_1_(^15^N) =124.39 ppm of **c)** 2D hNH spectra and **d)** expanded regions from the spectra in **c**. Spectra colored in blue were recorded at 850 MHz and spectra in red were measured at 1200 MHz.

We also recorded 2D hNH spectra on the ^13^C-^15^N labeled, and fully protonated, Rpo4/7 protein complex which is a subcomplex of archeal RNA polymerase II (Figure 8). Despite the full protonation, the spectra are nicely resolved, and show higher resolution at 1200 MHz. Peak picking performed in the amide region of the 2D spectra shows a clear increase in the number of picked peaks (using CCPNmr) from 80 to 98 peaks when comparing 850 MHz to 1200 MHz (Figure S6).

**Figure 8:**
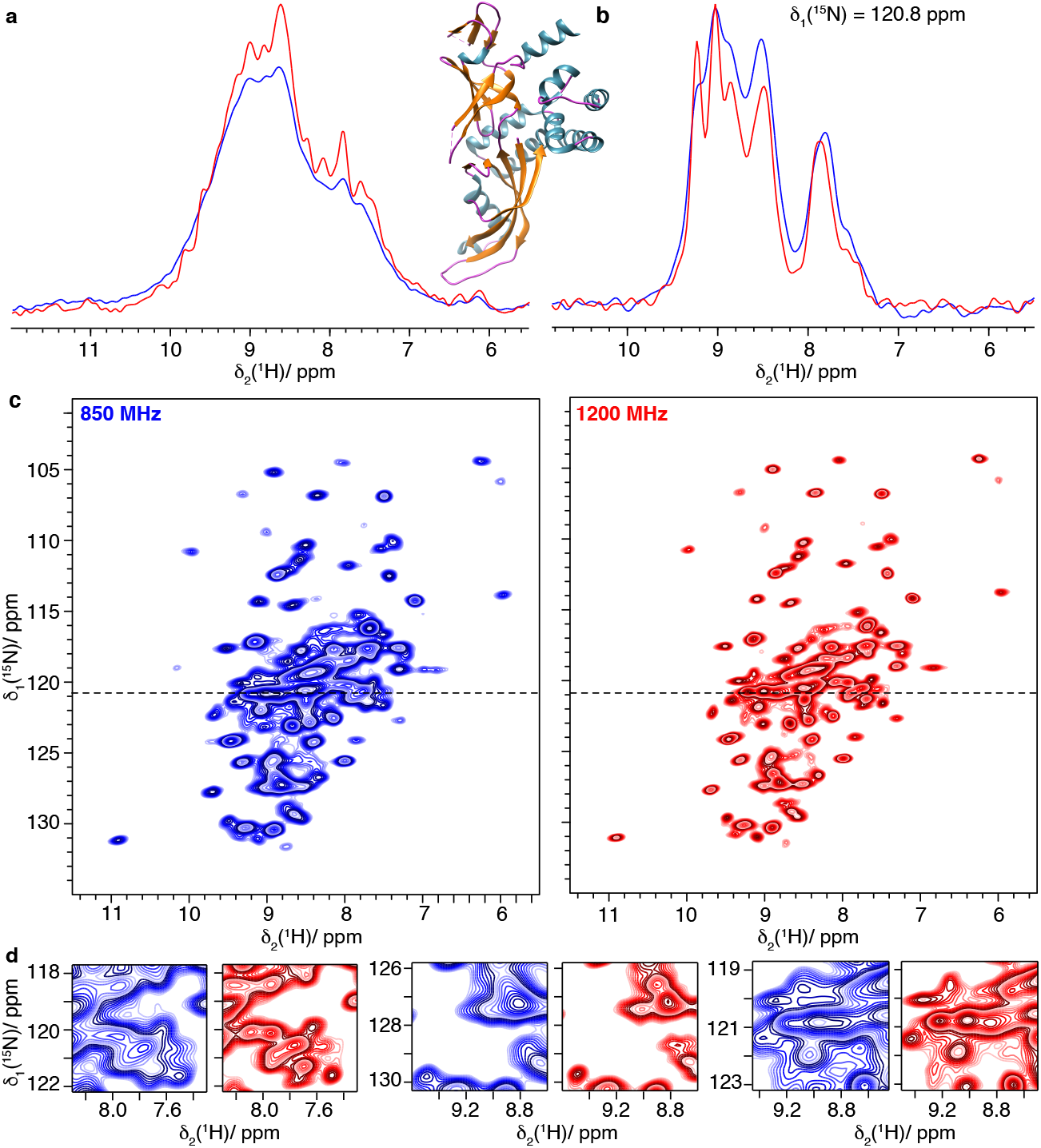
The Rpo4/7 protein complex (Rpo4C36S/Rpo7K123C). **a)** 1D-hnH spectra and structural model of Rpo4/7 (PDB ID: 1GO3) (Todone et al. 2001), **b)** one-dimensional trace at δ_1_(^15^N)=120.8 ppm of **c)** 2D hNH spectra and **d)** expanded regions from the spectra in **c**. Spectra colored in blue were recorded at 850 MHz and spectra in red were measured at 1200 MHz.

Furthermore, ^1^H-detected 2D spectra were acquired on fully protonated and ^13^C-^15^N labeled ASC filaments (Figure 9). The six *α*-helices forming the monomer (Liepinsh et al. 2003; Sborgi et al. 2015; Ravotti et al. 2016) make it more challenging for solid-state NMR due to the narrower distribution of chemical shifts and broader lines (due to stronger dipolar couplings in *α*-helices) (Malär et al. 2019) compared to proteins with higher variety in secondary structure. The higher magnetic field brings improved resolution in 2D hNH spectra (Figure 9a) which is again documented by more peaks picked automatically (142 and 170 peaks in hNH spectrum at 850 MHz and 1200 MHz, respectively, Figure S7).

**Figure 9:**
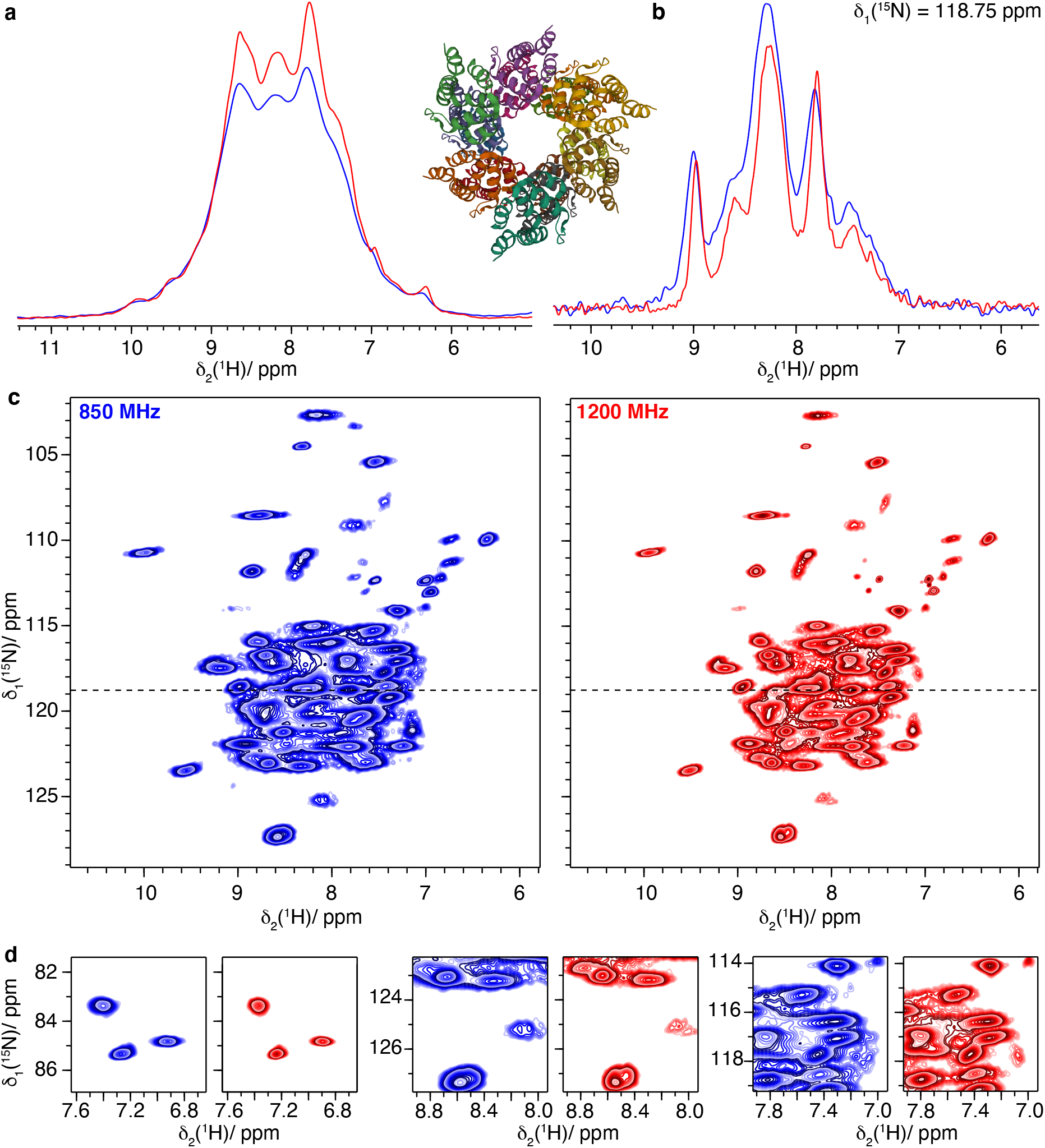
The filaments of PYRIN domain of mouse ASC. **a)** 1D-hnH spectra and structural model of ASC filaments (PDB ID: 2N1F) (Sborgi et al. 2015), **b)** one-dimensional trace at δ_1_(^15^N)=118.75 ppm of **c)** 2D hNH spectra and **d)** expanded regions from the spectra in **c**. Spectra colored in blue were recorded at 850 MHz and spectra in red were measured at 1200 MHz.

### ^1^H-detected ^1^H-^13^C correlation spectroscopy

A major limitation of proton-detected solid-state NMR spectroscopy today is the difficulty to observe H*α* and side-chain protons. First, deuteration is often used to improve resolution, so that solely H_N_ are present in the sample. Detection of H*α* and side-chain protons requires fully protonated samples, with the concomitant increase in line widths, resulting in poorer sensitivity and resolution. While H*α* and CH_3_ could often be detected already at lower fields, and even resolved in smaller proteins, CH_2_ remained in many cases severely broadened, as well as poorly dispersed due to the often highly similar chemical shifts of CH_2_ protons (Struppe et al. 2017). To illustrate the benefits of higher magnetic fields on protonated samples in this context, we recorded hCH spectra on three protein systems: ASC, Rpo4/7 and HET-s(218-289). The spectra are shown in Figures 10-12. For ASC and Rpo4/7, the 1200 MHz spectra are compared to spectra recorded at 850 MHz in Figures 10 and 11 panels c-e. The traces shown in Figure 10b and Figure 11b illustrate the gain in resolution for CH_2_ protons, here likely glycine CH_2_ groups. At higher field, a substantial gain in sensitivity and resolution is observed (Figure 10a,11a) in the hCH spectra. The improvements in resolution are clearly visible for the 2D hCH spectra in Figure 10c, d and 11c, d (compare respective expanded regions and trace along F2 in Figure 10b,11b), and especially for the CH_2_ resonance region shown in Figure 10c, 11c.

**Figure 10:**
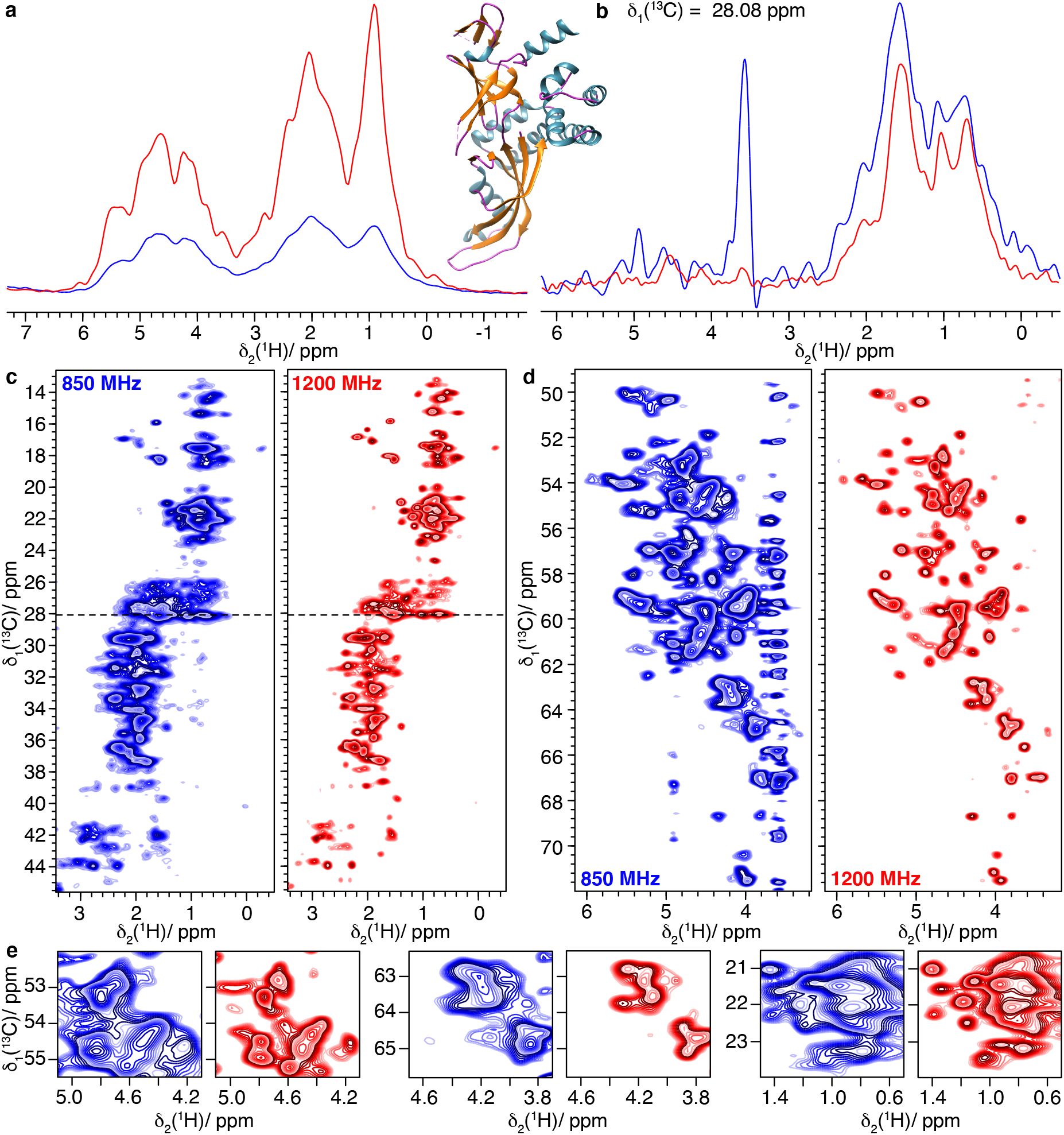
The Rpo4/7 protein complex (Rpo4C36S/Rpo7K123C). **a)** 1D-hcH spectra and structural model of Rpo4/7 (PDB 1GO3) (Todone et al. 2001), **b)** one-dimensional trace at δ_1_(^13^C)=28.08 ppm of **c)-d)** 2D hCH spectra and **e)** expanded regions from the spectra in **c-d**. Spectra colored in blue were recorded at 850 MHz and spectra in red were measured at 1200 MHz.

**Figure 11:**
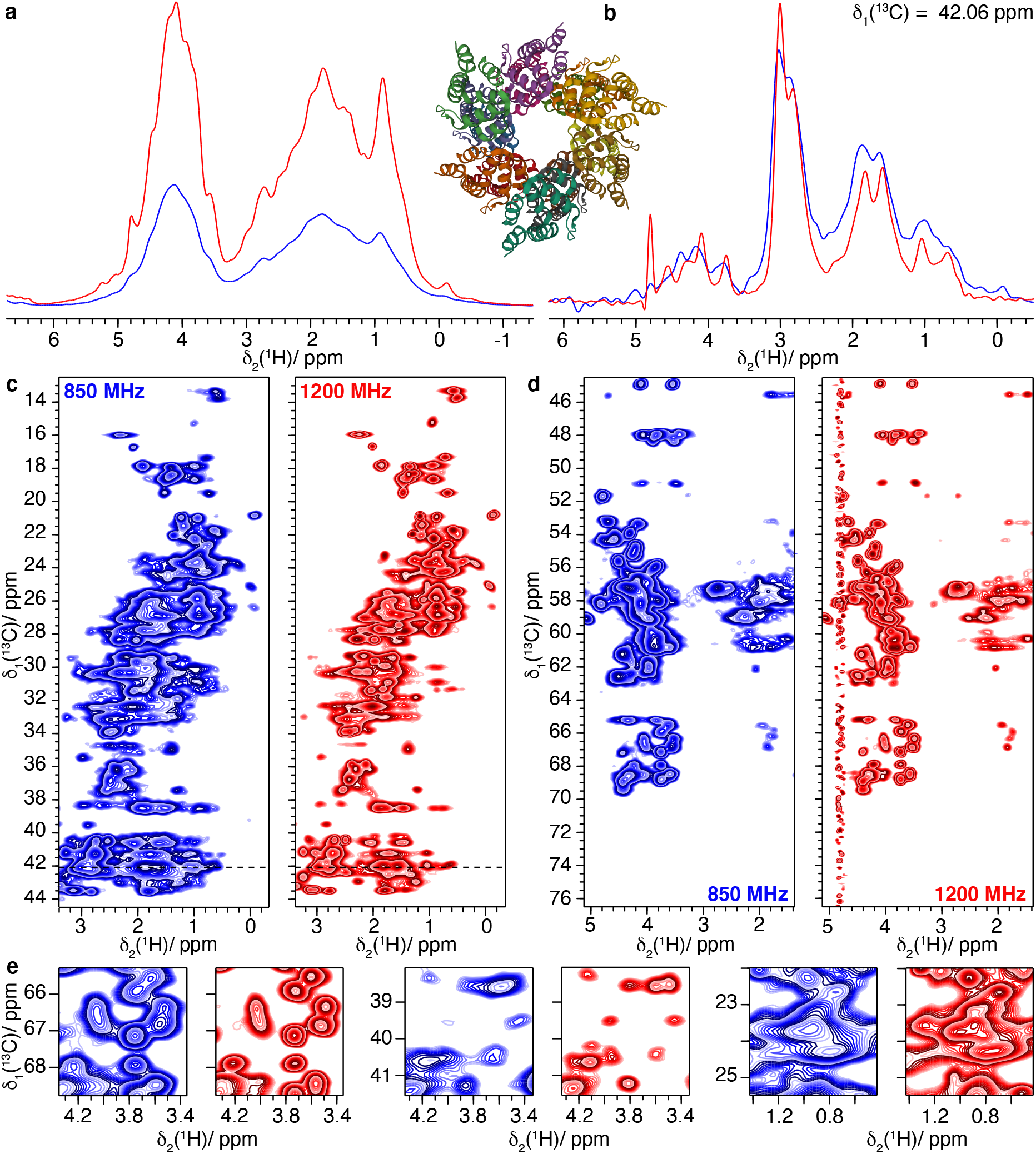
The filaments of PYRIN domain of mouse ASC. **a)** 1D-hcH spectra and structural model of ASC filame,ts (PDB 2N1F) (Sborgi et al. 2015), **b)** one-dimensional trace at δ_1_(^13^C)=42.06 ppm of **c)-d)** 2D hCH spectra and **e)** expanded regions from the spectra in **c-d**. Spectra colored in blue were recorded at 850 MHz and spectra in red were measured at 1200 MHz.

**Figure 12:**
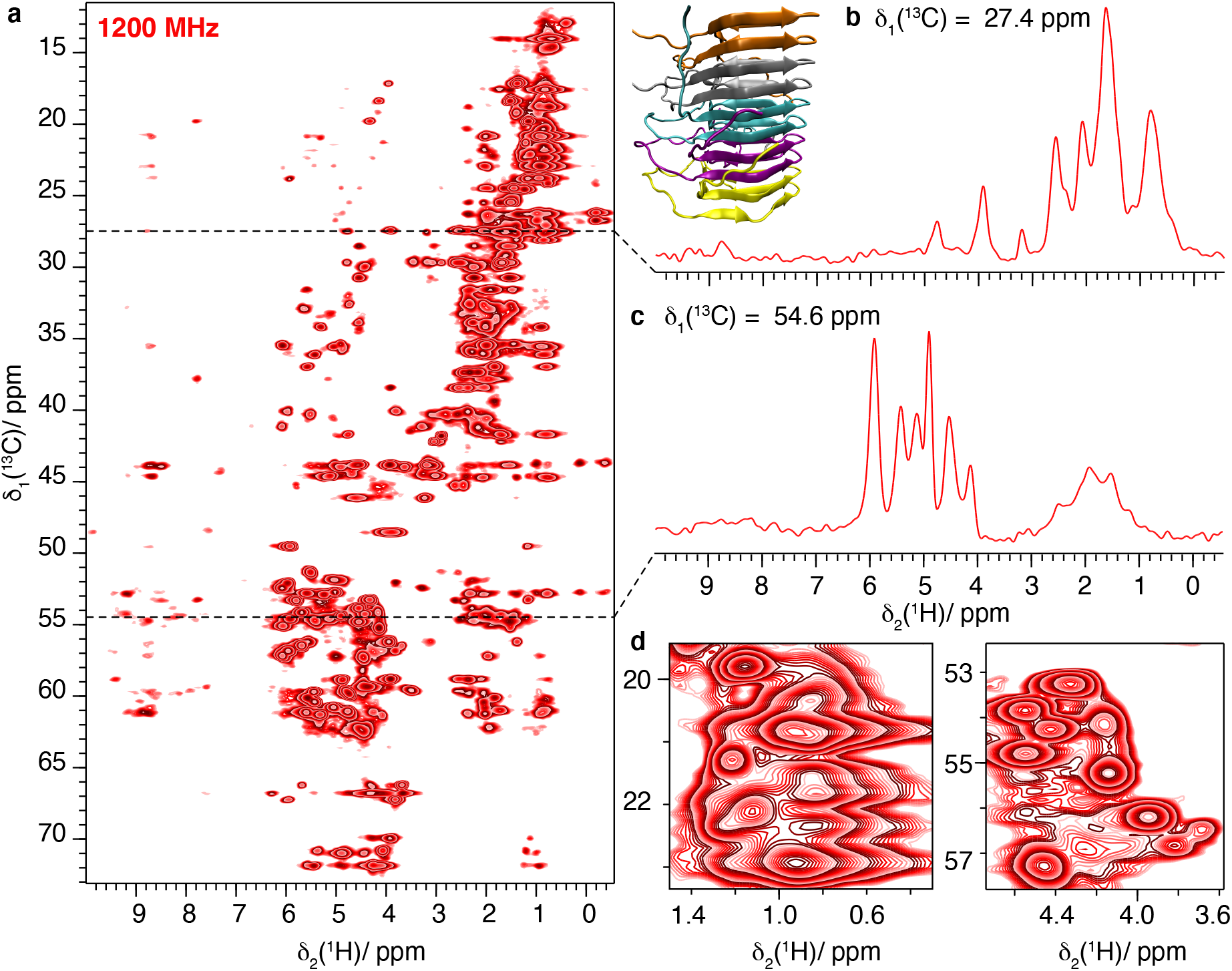
HET-s(218-289) fibrils fully protonated at 1200 MHz and 100 kHz MAS **a)** 1D-hCH spectrum and structural model of HET-s(218-289)(PDB ID: 2RNM)(Wasmer et al. 2008), **b)** one-dimensional trace at δ_1_(^13^C)=118.75 ppm, **c)** at 54.6 ppm and **d)** expanded regions from the spectrum in **a**.

For HET-s, only a 1200 MHz spectrum was recorded, and is shown in Figure 12. The excellent SNR obtained on this small protein allows even to detect further correlations than only one-bond. Indeed, entire spin systems can be observed for several amino acids. The spin diffusion under spin-lock condition during the CP actually seems to provide efficient polarization transfer, despite the scaling of the homonuclear dipolar couplings by a factor of −0.5 (Rhim, Pines, and Waugh 1970). This is due to the absence of chemical-shift differences in the rotating frame leading to a strong-coupling situation for the homonuclear dipolar couplings, corresponding to the laboratory frame situation for almost degenerate chemical shifts. (Xue et al. 2018)

In order to quantify the improvement in resolution when going to higher field, we compared the linewidths of ten randomly selected isolated peaks at 850 and 1200 MHz (Δ_ppm_(1200)/ Δ_ppm_(850), Figure S8). We observe that the improvement on the ppm scale is of about the expected value of 0.71 obtained from the field ratio. In the 2D hCH spectra recorded on the fully protonated samples a larger improvement is achieved, due to the more pronounced narrowing effect from the increased chemical-shift separation at higher field. The measurement of the bulk amide proton T_2_’(^1^H_N_) relaxation times (in Table S5) supports systematically the presence of this effect for all samples.

Access to information on side-chains is very important in sequential assignments, since notably the ^13^C shifts allow to identify amino-acid types. These frequencies were however difficult to observe when ^1^H-detection is used, since polarization transfer had to start on the H_N_, to go all the way out to side-chains carbons, and back to be detected. In protonated systems, CH_2_ were often too broad to be of use; this clearly is no longer the case at highest available fields and fastest spinning frequencies. 3D versions of these spectra e.g. hCCH spectra, shall thus allow to consider assignments of the aliphatic resonances at high field.

Distances between side-chain atoms are also very welcome as structural restraints; the better resolution can add CH and CH_2_ groups to the already used CH_3_ protons in selectively labeled samples (Agarwal et al. 2014; Agarwal and Reif 2008; Xue et al. 2018). Interestingly, the better resolution also paves the way for the detailed analysis of side-chain dynamics; typically, ^13^C relaxation measurements become possible above 60 kHz MAS (Smith et al. 2016) because ^13^C spin-diffusion is sufficiently suppressed. However, ^13^C detection is arduous at these MAS frequencies, and proton detection delivers better sensitivity. The increased resolution in the ^1^H dimension, at higher field, is therefore an important advantage in this context.

## Conclusions

We herein presented our first protein solid-state NMR spectra recorded at 1200 MHz revealing a significant gain in sensitivity and resolution for a variety of protein samples, ranging from amyloid fibrils, viral capsid proteins, protein complexes to helicases using ^13^C-, as well as ^1^H-detected experiments. For the samples described here, the improvement in resolution is variable but present for all samples.

The gain in resolution for ^13^C-detected spectra of large proteins will push the current resonance assignment limitations due to reduced spectral overlap and thereby provides an alternative to laborious biochemical approaches, for example segmental isotope labelling, or time-consuming spectroscopic methods such as 4D and 5D spectra. It will further allow studying structural and dynamic changes of high-molecular weight proteins upon interaction with other proteins, nucleic acids or small-molecule drugs, since the fate of more isolated peaks can be followed.

For proton-detected spectra, we observe an increase in resolution resulting from the increased chemical-shift dispersion as well as the reduced coherent contribution to the line width at higher magnetic field. This was shown to be stronger for protonated samples, due to their denser proton dipolar network. The high field thus allows to resolve aliphatic side-chain resonances and to characterize their structural properties as well as the dynamics. Importantly, the improved resolution and the concomitant gain in sensitivity at 1200 MHz, notably for the CH_2_ and CH_3_ groups, creates several new spectroscopic opportunities. First, side-chain resonances central in amino-acid identification and sequential assignments become more conveniently accessible. Second, the measurement of distance restraints involving sidechain atoms comes into reach also for uniformly labeled samples. Last but not least, it renders the investigation of sidechain dynamics via ^13^C relaxation a realistic objective.

## Materials and methods

### Sample preparation

^13^C-^15^N labeled protein samples were prepared as described in the literature: HET-s(218-289) fibrils (van Melckebeke et al. 2010), DnaB complexed with ADP:AlF_4_^-^ and DNA (Wiegand et al. 2019), Rpo4/7 protein complex of two subunits of RNA polymerase II (Torosyan et al. 2019) and filaments of PYRIN domain of mouse apoptosis-associated speck-like (ASC) protein containing a caspase-recruitment domain (Ravotti et al. 2016; Sborgi et al. 2015). The detailed protocols for TmcA, type 1 pili and ACNDVc will be described in forthcoming publications. ^2^H-^13^C-^15^N labeled and 100 % re-protonated Hepatitis B Virus Capsid (dCp149) was prepared as described by (Lecoq et al. 2019) while ^2^H-^13^C-^15^N labeled NS4B (dNS4B) was synthesized in H_2_O (Jirasko et al. 2020)).

### ^13^C-dectected spectroscopy

Solid-state NMR spectra were acquired on a wide-bore 850 MHz Bruker Avance III and on a standard-bore 1200 MHz Bruker Avance NEO spectrometer. ^13^C-detected solid-state NMR spectra were recorded using 3.2 mm Bruker Biospin “E-free probes”. The MAS frequency was set to 17.0 and 20.0 kHz at 850 MHz and 1200 MHz, respectively. The sample temperature was set to 278 K using the water line as an internal reference (Böckmann et al. 2009). The 2D spectra were processed with the software TOPSPIN (version 3.5 and 4.0.6, Bruker Biospin) with a shifted (3.0) squared cosine apodization function and automated baseline correction in the indirect and direct dimension. For further experimental details see Table S1.

### ^1^H-detected spectroscopy

The ^1^H detected spectra were acquired at 100 kHz MAS frequency using a Bruker 0.7 mm triple-resonance probe. The magic angle has been adjusted “on sample” by measuring *T*_2_’ proton transverse relaxation times and adjustment of the magic angle until the longest relaxation times were obtained (see Figure S2). The sample temperature was set to 293 K as determined from the supernatant water resonance (Böckmann et al. 2009; Gottlieb, Kotlyar, and Nudelman 1997). Two-dimensional (2D) fingerprint spectra (hNH) were recorded on dCp149, dNS4B Rpo4/7 protein complex and ASC, and 2D-hCH spectra on the Rpo4/7 protein complex, ASC filaments and HET-s(218-289). At both spectrometer frequencies, the 2D spectra were recorded with identical acquisition parameters for each protein sample (see Tables S2 to S4). All ^1^H detected spectra were processed using Topspin 4.0.6 (Bruker Topspin) with zero filling to the double amount of data points and a shifted sine-bell apodization function in direct and indirect dimensions with SSB=2.5. The direct dimension was truncated to 12.9 ms during processing of measurements at both magnetic field strengths. Spectral analysis was performed using CcpNmr Anlaysis 2.4.2 (Fogh et al. 2002; Stevens et al. 2011; Vranken et al. 2005). The spectra were referenced to 4,4-dimethyl-4-silapentane-1-sulfonic acid (DSS). One-dimensional spectra were scaled to the same noise level and 2D spectra to the same intensity level to compare spectra recorded at the two magnetic fields.

## Acknowledgements

Financial support by the ETH Zurich, the Department of Chemistry and Applied Biosciences, and the ETH foundation has been essential for obtaining the spectrometer. The scientific projects featured are supported by an ERC Advanced Grant (B.H.M., grant number 741863, Faster), by the Swiss National Science Foundation (B.H.M., grant number 200020_159707 and 200020_188711), an ETH Research Grant ETH-43 17-2 (T.W.), the French Agence Nationale de Recherches sur le Sida et les hépatites virales (ANRS, ECTZ71388 & ECTZ100488), the CNRS (CNRS-Momentum 2018), the LABEX ECOFECT (ANR-11-LABX-0048) within the Université de Lyon program Investissements d’Avenir (ANR-11-IDEX-0007). We thank Dr. Patrick Wikus and Dr. Rainer Kümmerle from Bruker Schweiz AG for their support.

## References

Abragam, A. 1961. The Principles of Nuclear Magnetism. Oxford Science Publications.

Agarwal, Vipin, Susanne Penzel, Kathrin Székely, Riccardo Cadalbert, Emilie Testori, Andres Oss, Jaan Past, et al. 2014. “De Novo 3D Structure Determination from Sub-Milligram Protein Samples by Solid-State 100 KHz MAS NMR Spectroscopy.” Angewandte Chemie, International Edition in English 53 (45): 12253–56. https://doi.org/10.1002/anie.201405730.

Agarwal, Vipin, and Bernd Reif. 2008. “Residual Methyl Protonation in Perdeuterated Proteins for Multi-Dimensional Correlation Experiments in MAS Solid-State NMR Spectroscopy.” Journal Of Magnetic Resonance 194 (1): 16–24. https://doi.org/10.1016/j.jmr.2008.05.021.

Andreas, Loren B, Tanguy Le Marchand, Kristaps Jaudzems, and Guido Pintacuda. 2015. “High-Resolution Proton-Detected NMR of Proteins at Very Fast MAS.” Journal Of Magnetic Resonance 253 (C): 36–49. https://doi.org/10.1016/j.jmr.2015.01.003.

Aue, WP, E Bartholdi, and RR Ernst. 1976. “Two-Dimensional Spectroscopy. Application to Nuclear Magnetic Resonance.” The Journal of Chemical Physics 64: 2229.

Barbet-Massin, Emeline, Andrew J Pell, Joren S Retel, Loren B Andreas, Kristaps Jaudzems, W Trent Franks, Andrew J Nieuwkoop, et al. 2014. “Rapid Proton-Detected NMR Assignment for Proteins with Fast Magic Angle Spinning.” Journal Of The American Chemical Society 136 (35): 12489–97. https://doi.org/10.1021/ja507382j.

Bazin, Alexandre, Mickaël V Cherrier, Irina Gutsche, Joanna Timmins, and Laurent Terradot. 2015. “Structure and Primase-Mediated Activation of a Bacterial Dodecameric Replicative Helicase.” Nucleic Acids Research 43 (17): 8564–76. https://doi.org/10.1093/nar/gkv792.

Böckmann, Anja, Matthias Ernst, and Beat H Meier. 2015. “Spinning Proteins, the Faster, the Better?” Journal Of Magnetic Resonance 253 (C): 71–79. https://doi.org/10.1016/j.jmr.2015.01.012.

Böckmann, Anja, Carole Gardiennet, Rene Verel, Andreas Hunkeler, Antoine Loquet, Guido Pintacuda, Lyndon Emsley, Beat H Meier, and Anne Lesage. 2009. “Characterization of Different Water Pools in Solid-State NMR Protein Samples.” Journal of Biomolecular NMR 45 (3): 319–27. https://doi.org/10.1007/s10858-009-9374-3.

Castellani, F, B van Rossum, A Diehl, M Schubert, K Rehbein, and Hartmut Oschkinat. 2002. “Structure of a Protein Determined by Solid-State Magic-Angle-Spinning NMR Spectroscopy.” Nature 420 (6911): 98–102.

Chimnaronk, Sarin, Tateki Suzuki, Tetsuhiro Manita, Yoshiho Ikeuchi, Min Yao, Tsutomu Suzuki, and Isao Tanaka. 2009. “RNA Helicase Module in an Acetyltransferase That Modifies a Specific TRNA Anticodon.” The EMBO Journal 28 (9): 1362–73. https://doi.org/10.1038/emboj.2009.69.

Colombo, M G, Beat H Meier, and RR Ernst. 1988. “Rotor-Driven Spin Diffusion in Natural-Abundance 13-C Spin Systems.” Chemical Physics Letters 146 (3): 189.

Ernst, RR, and WA Anderson. 1965. “Application of Fourier Transform Spectrsocopy to Magnetic Resonance.” Review Of Scientific Instruments 37 (1): 93–102.

Fiaux, Jocelyne, Eric B. Bertelsen, Arthur L. Horwich, and Kurt Wüthrich. 2002. “NMR Analysis of a 900K GroEL GroES Complex.” Nature 418 (6894): 207–11. https://doi.org/10.1038/nature00860.

Fogh, Rasmus, John Ionides, Eldon Ulrich, Wayne Boucher, Wim Vranken, Jens P Linge, Michael Habeck, et al. 2002. “The CCPN Project: An Interim Report on a Data Model for the NMR Community.” Nature Structural Biology 9 (6): 416–18. https://doi.org/10.1038/nsb0602-416.

Gardiennet, Carole, Anne K Schütz, Andreas Hunkeler, Britta Kunert, Laurent Terradot, Anja Böckmann, and Beat H Meier. 2012. “A Sedimented Sample of a 59 KDa Dodecameric Helicase Yields High-Resolution Solid-State NMR Spectra.” Angewandte Chemie, International Edition in English 51 (31): 7855–58. https://doi.org/10.1002/anie.201200779.

Gor’kov, Peter L, Raiker Witter, Eduard Y Chekmenev, Farhod Nozirov, Riqiang Fu, and William W Brey. 2007. “Low-E Probe for (19)F-(1)H NMR of Dilute Biological Solids.” Journal Of Magnetic Resonance 189 (2): 182–89. https://doi.org/10.1016/j.jmr.2007.09.008.

Gottlieb, Hugo E, Vadim Kotlyar, and Abraham Nudelman. 1997. “NMR Chemical Shifts of Common Laboratory Solvents as Trace Impurities.” The Journal of Organic Chemistry 62 (21): 7512–15.

Gouttenoire, Jérôme, Roland Montserret, David Paul, Rosa Castillo, Simon Meister, Ralf Bartenschlager, Francois Penin, and Darius Moradpour. 2014. “Aminoterminal Amphipathic α-Helix AH1 of Hepatitis c Virus Nonstructural Protein 4B Possesses a Dual Role in RNA Replication and Virus Production.” PLoS Pathogens 10 (11): e1004501–17. https://doi.org/10.1371/journal.ppat.1004501.

Habenstein, Birgit, Antoine Loquet, Songhwan Hwang, Karin Giller, Suresh Kumar Vasa, Stefan Becker, Michael Habeck, and Adam Lange. 2015. “Hybrid Structure of the Type 1 Pilus of Uropathogenic Escherichia Coli.” Angewandte Chemie (International Ed. in English) 54 (40): 11691–95. https://doi.org/10.1002/anie.201505065.

Hahn, Erik, Peter Wild, Uta Hermanns, Peter Sebbel, Rudi Glockshuber, Marcus Häner, Nicole Taschner, Peter Burkhard, Ueli Aebi, and Shirley A. Müller. 2002. “Exploring the 3D Molecular Architecture of Escherichia Coli Type 1 Pili.” Journal of Molecular Biology 323 (5): 845–57. https://doi.org/10.1016/s0022-2836(02)01005-7.

Ikeuchi, Yoshiho, Kei Kitahara, and Tsutomu Suzuki. 2008. “The RNA Acetyltransferase Driven by ATP Hydrolysis Synthesizes N4-Acetylcytidine of TRNA Anticodon.” The EMBO Journal 27 (16): 2194–2203. https://doi.org/10.1038/emboj.2008.154.

Jirasko, Vlastimil, Nils-Alexander Lakomek, Susanne Penzel, Marie-Laure Fogeron, Ralf Bartenschlager, Beat H Meier, and Anja Böckmann. 2020. “Proton-Detected Solid-State NMR of the Cell-Free Synthesized α-Helical Transmembrane Protein NS4B from Hepatitis c Virus.” Chembiochem: A European Journal of Chemical Biology 21 (10): 1453–60. https://doi.org/10.1002/cbic.201900765.

Lauber, Chris, Stefan Seitz, Simone Mattei, Alexander Suh, Jürgen Beck, Jennifer Herstein, Jacob Börold, et al. 2017. “Deciphering the Origin and Evolution of Hepatitis b Viruses by Means of a Family of Non-Enveloped Fish Viruses.” Cell Host and Microbe 22 (3): 387–399.e6. https://doi.org/10.1016/j.chom.2017.07.019.

Lecoq, Lauriane, Maarten Schledorn, Shishan Wang, Susanne Smith-Penzel, Alexander A. Malär, Morgane Callon, Michael Nassal, Beat H. Meier, and Anja Böckmann. 2019. “100 KHz MAS Proton-Detected NMR Spectroscopy of Hepatitis B Virus Capsids.” Frontiers in Molecular Biosciences 6: 58. https://doi.org/10.3389/fmolb.2019.00058.

Lecoq, Lauriane, Shishan Wang, Thomas Wiegand, Stephane Bressanelli, Michael Nassal, Beat H Meier, and Anja Böckmann. 2018. “Localizing Conformational Hinges by NMR: Where Do Hepatitis b Virus Core Proteins Adapt for Capsid Assembly?” Chemphyschem 19 (11): 1336–40. https://doi.org/10.1002/cphc.201800211.

Liepinsh, Edvards, Raitis Barbals, Edgar Dahl, Anatoly Sharipo, Eike Staub, and Gottfried Otting. 2003. “The Death-Domain Fold of the ASC PYRIN Domain, Presenting a Basis for PYRIN/PYRIN Recognition.” Journal Of Molecular Biology 332 (5): 1155–63. https://doi.org/10.1016/j.jmb.2003.07.007.

Maeda, Hideaki, and Yoshinori Yanagisawa. 2019. “Future Prospects for NMR Magnets: A Perspective.” Journal of Magnetic Resonance (San Diego, Calif.: 1997) 306 (September): 80–85. https://doi.org/10.1016/j.jmr.2019.07.011.

Malär, Alexander A, Susanne Smith-Penzel, Gian-Marco Camenisch, Thomas Wiegand, Ago Samoson, Anja Böckmann, Matthias Ernst, and Beat H Meier. 2019. “Quantifying Proton NMR Coherent Linewidth in Proteins under Fast MAS Conditions: A Second Moment Approach.” Physical Chemistry Chemical Physics: PCCP 420: 98–16. https://doi.org/10.1039/C9CP03414E.

McDermott, A, T Polenova, A Bockmann, K W Zilm, E K Paulson, R W Martin, G T Montelione, and E K Paulsen. 2000. “Partial NMR Assignments for Uniformly (13C, 15N)-Enriched BPTI in the Solid State.” Journal of Biomolecular NMR 16 (3): 209–19.

Melckebeke, Hélène van, Christian Wasmer, Adam Lange, Eiso Ab, Antoine Loquet, Anja Böckmann, and Beat H Meier. 2010. “Atomic-Resolution Three-Dimensional Structure of HET-s(218-289) Amyloid Fibrils by Solid-State NMR Spectroscopy.” Journal Of The American Chemical Society 132 (39): 13765–75. https://doi.org/10.1021/ja104213j.

Nimerovsky, Evgeny, Kumar Tekwani Movellan, Xizhou Zhang, Marcel C. Forster, Eszter Najbauer, Kai Xue, Rıza Dervişoğlu, et al. 2021. “Proton Detected Solid-State NMR of Membrane Proteins at 28 Tesla and 100 Khz Magic-Angle Spinning,” March. https://www.preprints.org/manuscript/202103.0691/v1.

Penzel, Susanne, Andres Oss, Mai-Liis Org, Ago Samoson, Anja Böckmann, Matthias Ernst, and Beat H Meier. 2019. “Spinning Faster: Protein NMR at MAS Frequencies up to 126 KHz.” Journal of Biomolecular NMR 73 (1-2): 19–29. https://doi.org/10.1007/s10858-018-0219-9.

Pervushin, K, Roland Riek, G Wider, and K Wüthrich. 1997. “Attenuated T2 Relaxation by Mutual Cancellation of Dipole-Dipole Coupling and Chemical Shift Anisotropy Indicates an Avenue to NMR Structures of Very Large Biological Macromolecules in Solution.” Proceedings Of The National Academy Of Sciences Of The United States Of America 94 (23): 12366–71. https://doi.org/10.1073/pnas.94.23.12366.

Raleigh, DP, MH Levitt, and Robert G Griffin. 1988. “Rotational Resonance in Solid State NMR.” Chemical Physics Letters 146: 71.

Ravotti, Francesco, Lorenzo Sborgi, Riccardo Cadalbert, Matthias Huber, Adam Mazur, Petr Broz, Sebastian Hiller, Beat H Meier, and Anja Böckmann. 2016. “Sequence-Specific Solid-State NMR Assignments of the Mouse ASC PYRIN Domain in Its Filament Form.” Biomolecular NMR Assignments 10 (1): 107–15. https://doi.org/10.1007/s12104-015-9647-6.

Rhim, WK, A Pines, and JS Waugh. 1970. “Violation of the Spin-Temperature Hypothesis.” Physical Review Letters 25 (4): 218.

Rosenzweig, Rina, and Lewis E. Kay. 2014. “Bringing Dynamic Molecular Machines into Focus by Methyl-TROSY NMR.” Annual Review of Biochemistry 83: 291–315. https://doi.org/10.1146/annurev-biochem-060713-035829.

Sborgi, Lorenzo, Francesco Ravotti, Venkata P Dandey, Mathias S Dick, Adam Mazur, Sina Reckel, Mohamed Chami, et al. 2015. “Structure and Assembly of the Mouse ASC Inflammasome by Combined NMR Spectroscopy and Cryo-Electron Microscopy.” Proceedings of the National Academy of Sciences of the United States of America 112 (43): 13237–42. https://doi.org/10.1073/pnas.1507579112.

Schledorn, Maarten, Alexander Malär, Anahit Torosyan, Susanne Penzel, Daniel Klose, Andres Oss, Mai-Liis Org, et al. 2020. “Protein NMR Spectroscopy at 150 KHz Magic-Angle Spinning Continues to Improve Resolution and Mass Sensitivity.” Chembiochem: A European Journal of Chemical Biology, June, cbic.202000341-26. https://doi.org/10.1002/cbic.202000341.

Siemer, A, A Arnold, C Ritter, T Westfeld, M Ernst, Roland Riek, and Beat H Meier. 2006. “Observation of Highly Flexible Residues in Amyloid Fibrils of the HET-s Prion.” Journal Of The American Chemical Society 128 (40): 13224–28.

Smith, Albert A. 2018. “Correction to: Characterization of Fibril Dynamics on Three Timescales by Solid-State NMR.” Journal of Biomolecular NMR 70 (3): 203–203. https://doi.org/10.1007/s10858-018-0170-9.

Smith, Albert A, Francesco Ravotti, Emilie Testori, Riccardo Cadalbert, Matthias Ernst, Anja Böckmann, and Beat H Meier. 2017. “Partially-Deuterated Samples of HET-s(218-289) Fibrils: Assignment and Deuterium Isotope Effect.” Journal of Biomolecular NMR 67 (2): 109–19. https://doi.org/10.1007/s10858-016-0087-0.

Smith, Albert A, Emilie Testori, Riccardo Cadalbert, Beat H Meier, and Matthias Ernst. 2016. “Characterization of Fibril Dynamics on Three Timescales by Solid-State NMR.” Journal of Biomolecular NMR 65 (3-4): 171–91. https://doi.org/10.1007/s10858-016-0047-8.

Stevens, Tim J, Rasmus H Fogh, Wayne Boucher, Victoria A Higman, Frank Eisenmenger, Benjamin Bardiaux, Barth-Jan van Rossum, Hartmut Oschkinat, and Ernest D Laue. 2011. “A Software Framework for Analysing Solid-State MAS NMR Data.” Journal of Biomolecular NMR 51 (4): 437–47. https://doi.org/10.1007/s10858-011-9569-2.

Struppe, Jochem, Caitlin M Quinn, Manman Lu, Mingzhang Wang, Guangjin Hou, Xingyu Lu, Jodi Kraus, et al. 2017. “Expanding the Horizons for Structural Analysis of Fully Protonated Protein Assemblies by NMR Spectroscopy at MAS Frequencies above 100 KHz.” Solid State Nuclear Magnetic Resonance 87 (July): 117–25. https://doi.org/10.1016/j.ssnmr.2017.07.001.

Takegoshi, K, S Nakamura, and T Terao. 2001. “C-13-H-1 Dipolar-Assisted Rotational Resonance in Magic-Angle Spinning NMR.” Chemical Physics Letters 344 (5-6): 631–37.

Takegoshi, K, Shinji Nakamura, and Takehiko Terao. 2003. “13C-1H Dipolar-Driven 13C-13C Recoupling without 13C Rf Irradiation in Nuclear Magnetic Resonance of Rotating Solids.” The Journal of Chemical Physics 118 (5): 2325–41.

Todone, F., P. Brick, F. Werner, R. O. Weinzierl, and S. Onesti. 2001. “Structure of an Archaeal Homolog of the Eukaryotic RNA Polymerase II RPB4/RPB7 Complex.” Molecular Cell 8 (5): 1137–43. https://doi.org/10.1016/s1097-2765(01)00379-3.

Torosyan, Anahit, Thomas Wiegand, Maarten Schledorn, Daniel Klose, Peter Güntert, Anja Böckmann, and Beat H Meier. 2019. “Including Protons in Solid-State NMR Resonance Assignment and Secondary Structure Analysis: The Example of RNA Polymerase II Subunits Rpo4/7.” Frontiers in Molecular Biosciences 6: 100–108. https://doi.org/10.3389/fmolb.2019.00100.

Vranken, Wim F, Wayne Boucher, Tim J Stevens, Rasmus H Fogh, Anne Pajon, Miguel Llinas, Eldon L Ulrich, John L Markley, John Ionides, and Ernest D Laue. 2005. “The CCPN Data Model for NMR Spectroscopy: Development of a Software Pipeline.” Proteins-Structure Function And Bioinformatics 59 (4): 687–96. https://doi.org/10.1002/prot.20449.

Wasmer, Christian, Adam Lange, Hélène van Melckebeke, Ansgar B Siemer, Roland Riek, and Beat H Meier. 2008. “Amyloid Fibrils of the HET-s(218-289) Prion Form a Beta Solenoid with a Triangular Hydrophobic Core.” Science (New York, NY) 319 (5869): 1523–26. https://doi.org/10.1126/science.1151839.

Wiegand, Thomas. 2018. “Segmental Isotope Labelling and Solid-State NMR of a 12 × 59 KDa Motor Protein: Identification of Structural Variability.” Journal of Biomolecular NMR 0 (0): 0–0. https://doi.org/10.1007/s10858-018-0196-z.

Wiegand, Thomas, Riccardo Cadalbert, Denis Lacabanne, Joanna Timmins, Laurent Terradot, Anja Böckmann, and Beat H Meier. 2019. “The Conformational Changes Coupling ATP Hydrolysis and Translocation in a Bacterial DnaB Helicase.” Nature Communications 10 (1): 1–11. https://doi.org/10.1038/s41467-018-07968-3.

Wiegand, Thomas, Carole Gardiennet, Riccardo Cadalbert, Denis Lacabanne, Britta Kunert, Laurent Terradot, Anja Böckmann, and Beat H Meier. 2016. “Variability and Conservation of Structural Domains in Divide-and-Conquer Approaches.” Journal of Biomolecular NMR 65 (2): 79–86. https://doi.org/10.1007/s10858-016-0039-8.

Wiegand, Thomas, Carole Gardiennet, Francesco Ravotti, Alexandre Bazin, Britta Kunert, Denis Lacabanne, Riccardo Cadalbert, et al. 2016. “Solid-State NMR Sequential Assignments of the N-Terminal Domain of HpDnaB Helicase.” Biomolecular NMR Assignments 10 (1): 13–23. https://doi.org/10.1007/s12104-015-9629-8.

Williamson, Mp, Tf Havel, and K. Wuthrich. 1985. “Solution Conformation of Proteinase Inhibitor-Iia from Bull Seminal Plasma by H-1 Nuclear Magnetic-Resonance and Distance Geometry.” Journal of Molecular Biology 182 (2): 295–315. https://doi.org/10.1016/0022-2836(85)90347-X.

Wüthrich, K. 2003. “NMR Studies of Structure and Function of Biological Macromolecules (Nobel Lecture).” Journal of Biomolecular NMR 27 (1): 13–39.

Wynne, S A, R A Crowther, and A G Leslie. 1999. “The Crystal Structure of the Human Hepatitis B Virus Capsid.” Molecular Cell 3 (6): 771–80.

Xue, Kai, Riddhiman Sarkar, Carina Motz, Sam Asami, Venita Decker, Sebastian Wegner, Zdeněk Tošner, and Bernd Reif 2018. “Magic-Angle Spinning Frequencies beyond 300 KHz Are Necessary to Yield Maximum Sensitivity in Selectively Methyl Protonated Protein Samples in Solid-State NMR.” The Journal of Physical Chemistry C 122 (28): 1–6. https://doi.org/10.1021/acs.jpcc.8b05600.

Xue, Kai, Riddhiman Sarkar, Daniela Lalli, Benita Koch, Guido Pintacuda, Zdenek Tosner, and Bernd Reif. 2020. “Impact of Magnetic Field Strength on Resolution and Sensitivity of Proton Resonances in Biological Solids.” Journal of Physical Chemistry C 124 (41): 22631–37. https://doi.org/10.1021/acs.jpcc.0c05407.

