## Supplementary material for "Biomolecular solid-state NMR spectroscopy at highest field: the gain in resolution at 1200 MHz": SI1p2GHz31022021finalA.pdf

#### Supplementary Figures

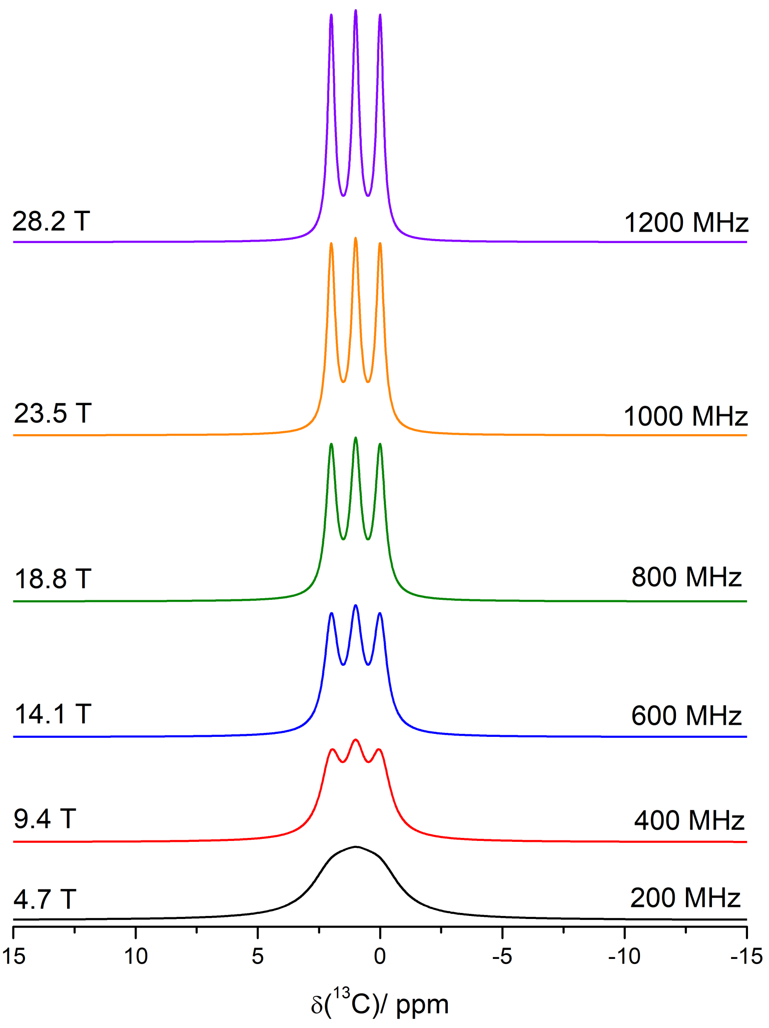


**Figure S1**: *NMR spectra gain in resolution on the ppm-scale by increasing the external magnetic field strength.* Simulated NMR spectra at different magnetic fields of three resonances assuming a constant full-width at half maximum (FWHM= 100 Hz). The integral of the resonances on the ppm-scale is kept constant.

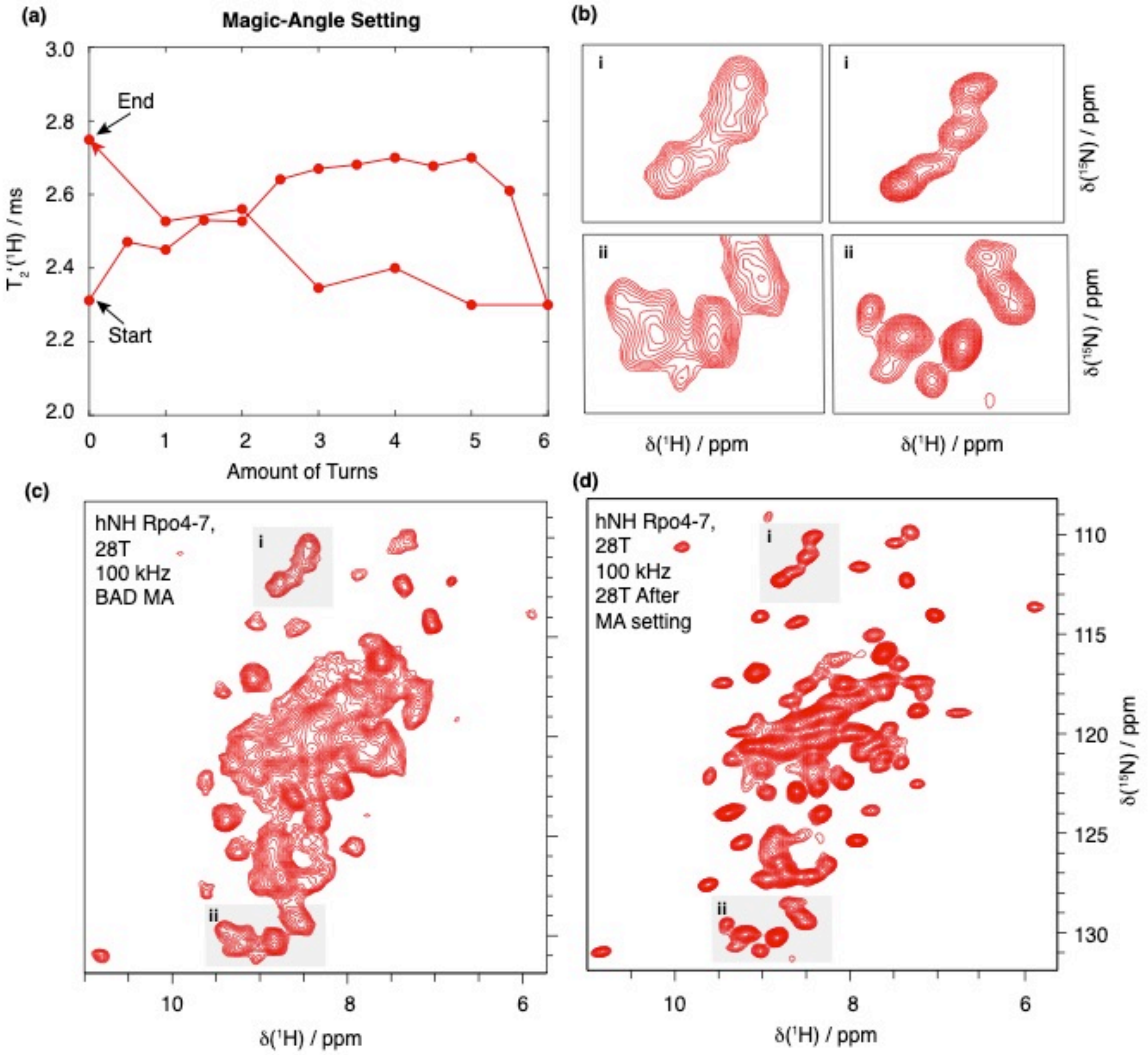


**Figure S2**: *"On sample" magic-angle setup by optimizing on the longest proton $T_2$' transverse relaxation times* **a** $T_2$'($^1H_N$) transverse relaxation time as function of the number of turns of the magic-angle screw on a nstandard bore 0.7 mm probe-head. **b** zooms of the 2D hNH spectra of Rpo4/7 protein complex recorded **c** before and **d** after magic angle optimization with $T_2$'-on-sample method.

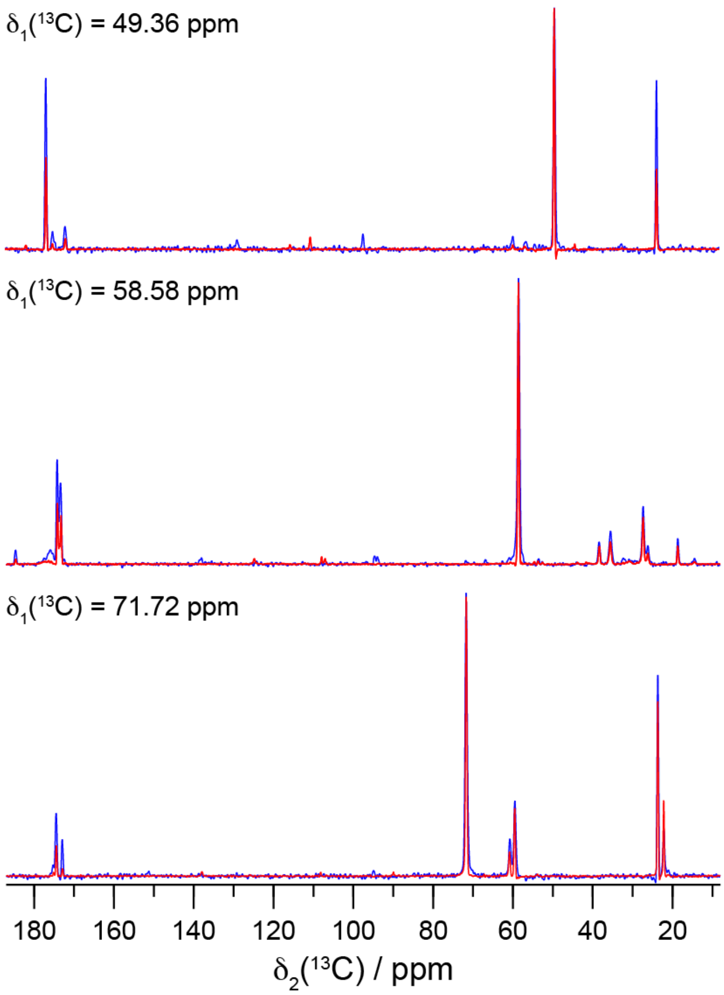


**Figure S3:** *The cross-peak intensities in 20 ms $^{13}C$-$^{13}C$ DARR is less at 1200 MHz compared to 850 MHz.* Three representative 1D traces along F2 taken from the DARR spectrum of HET-s(218-289) fibirls (Figure 2). The blue spectrum has been recorded at 850 MHz, the red spectrum at 1200 MHz. The spectra were scaled to the diagonal peak. In all cases, the cross-peak intensity at 1200 MHz t is lower than at 850 MHz.

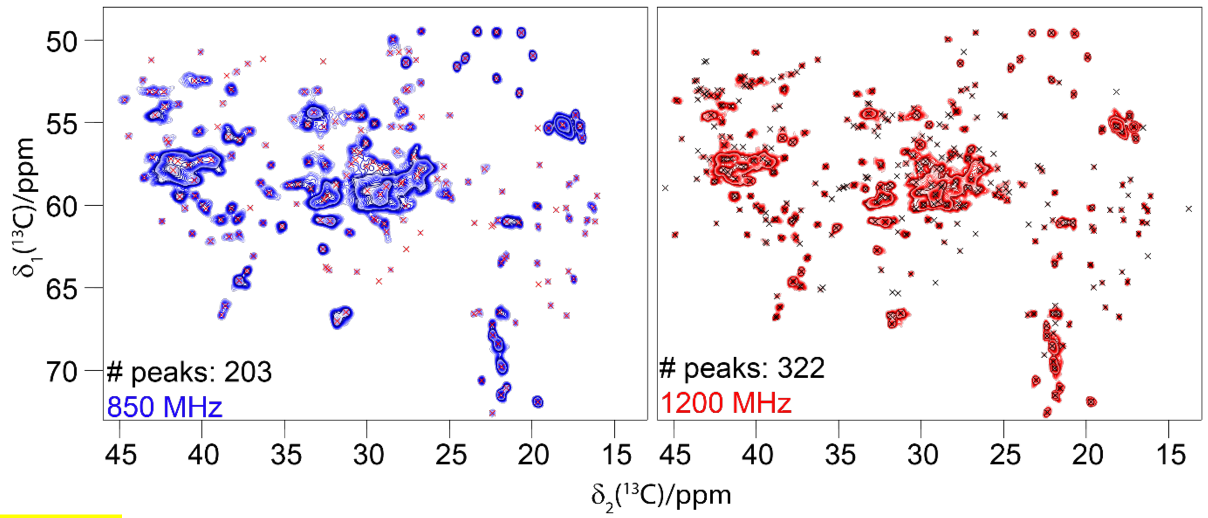


**Figure S4**: *The number of automatically picked peaks increases at higher magnetic-field strength.* Automatically picked resonances for DnaB (crosses) plotted on $^{13}C$-$^{13}C$ 20 ms DARR spectra recorded at 850 MHz (203 peaks) and 1200 MHz (322 peaks).

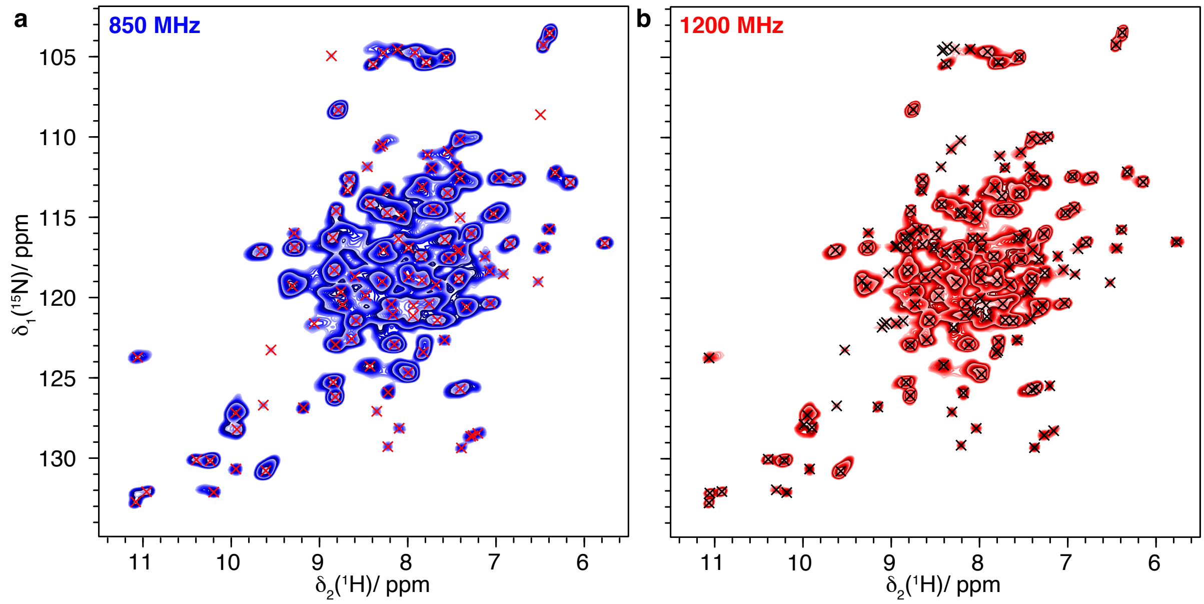


**Figure S5**: *The number of automatically picked peaks increases at higher magnetic-field strength.* Automatically picked resonances for dCp149 (crosses) plotted on 2D hNH spectra recorded at **a** 850 MHz (110 peaks) and **b** 1200 MHz (157 peaks). Note that number of peaks picked in the 1200 MHz spectrum is higher than the number of amino-acids in the protein due to the presence of four different molecules in the asymmetric unit of the T=4 icosahedral HBV capsid that causes peak splitting of Cp149 [1].

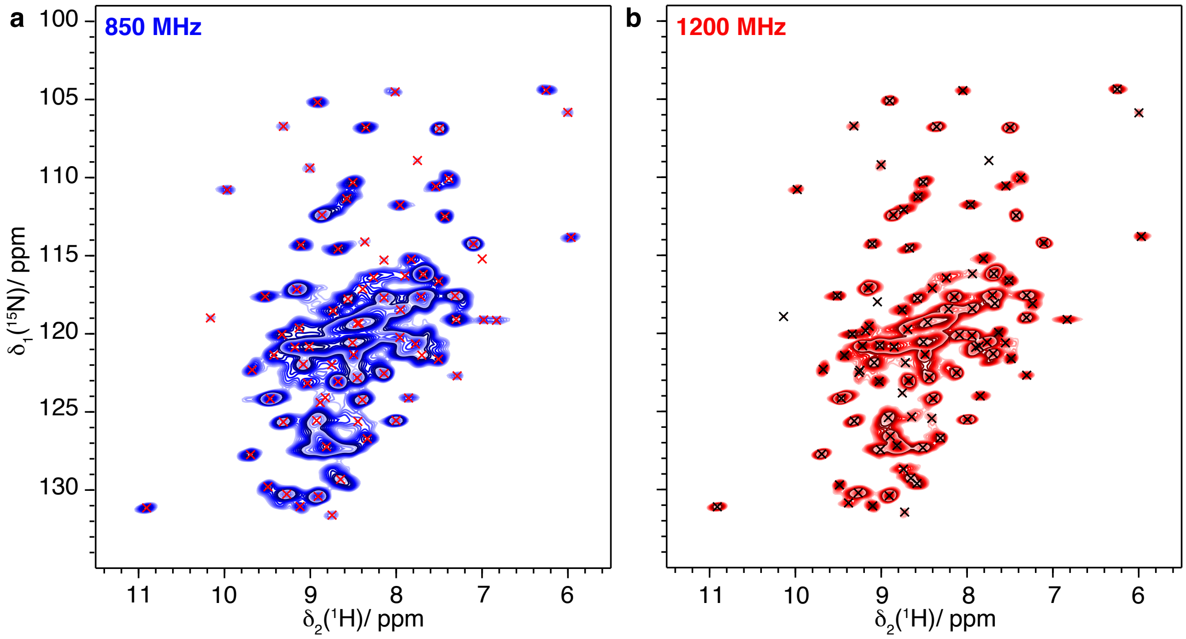


**Figure S6**: *The number of automatically picked peaks increases at higher magnetic-field strength.* Automatically picked resonances for the Rpo4/7 protein complex (Rpo4C36S/Rpo7K123C) (crosses) plotted on 2D hNH spectra recorded at **a** 850 MHz (80 peaks) and **b** 1200 MHz (98 peaks).

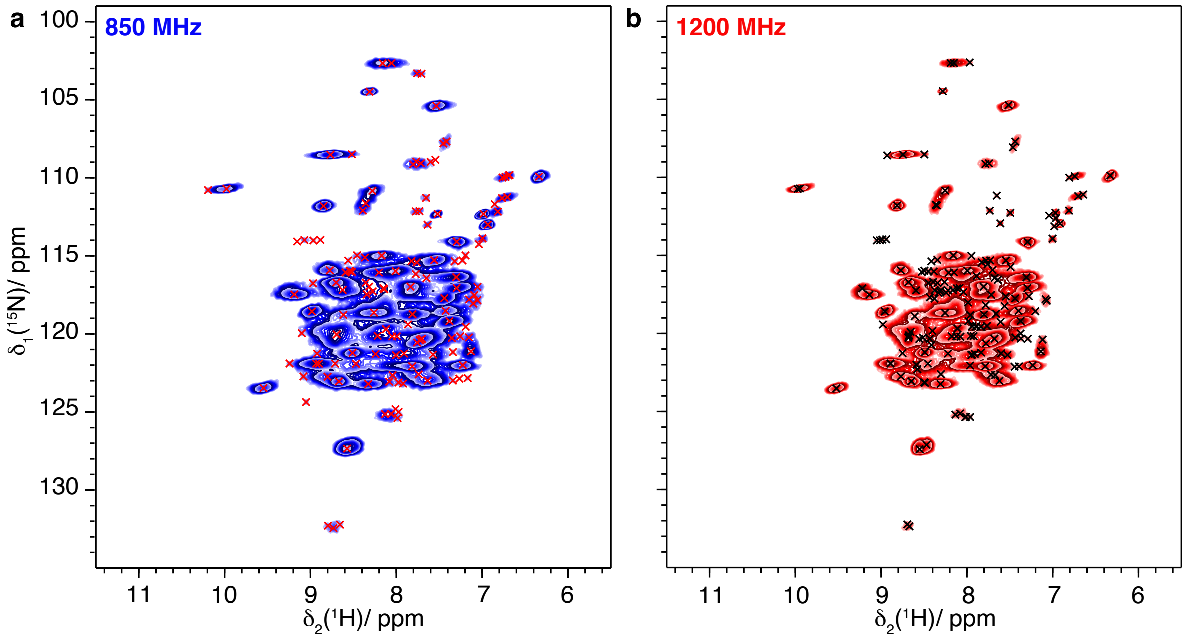


**Figure S7**: *The number of automatically picked peaks increases at higher magnetic-field strength.* Automatically picked resonances for the filaments of PYRIN domain of mouse ASC (crosses) plotted on 2D hNH spectra recorded at **a** 850 MHz (142 peaks) and **b** 1200 MHz (170 peaks).

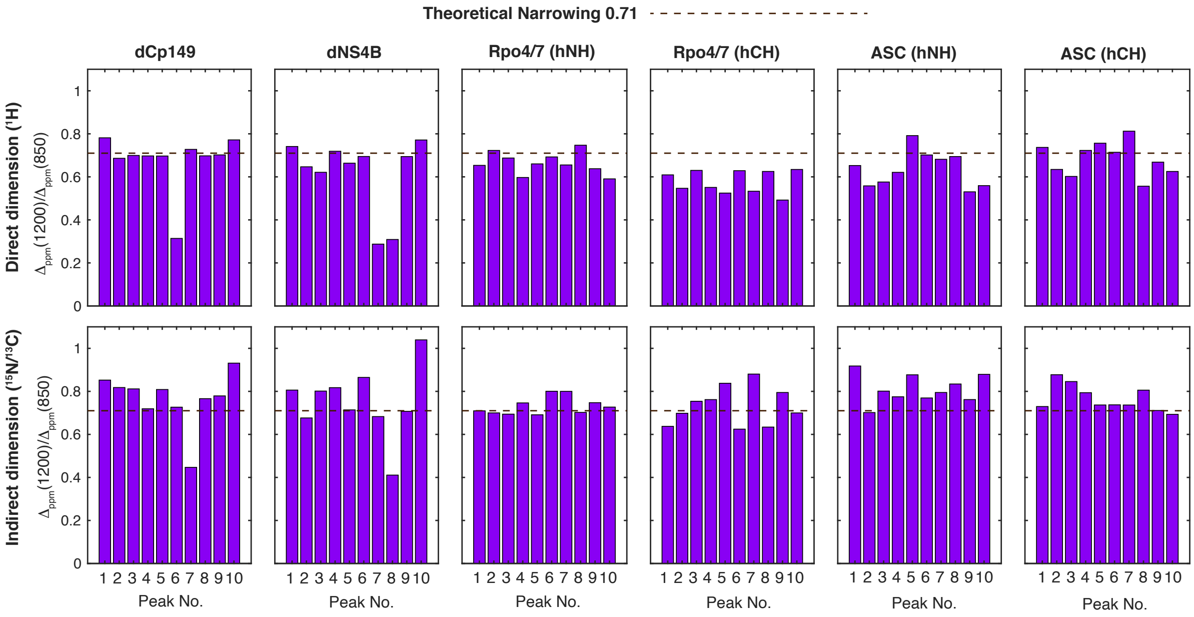


**Figure S8**: Comparison of the total linewidth ($\Delta_{ppm}(1200)$ / $\Delta_{ppm}(850)$) in ppm for both direct and indirect dimensions, for a set of ten isolated and randomly selected peaks for the five different protein systems. The theoretical narrowing value (dashed black line) is obtained by calculating the field ratio 850/1200 = 0.71.

**Supplementary table 1**: NMR parameters of 2D $^{13}C$-$^{13}C$ DARR correlation spectra recorded at 850 MHz (20.0 T) and 1200 MHz (28.2 T).

| **Experiment** | **DARR (DnaB)** | | | **DARR (HET-s(218-289))** | |
|---|---|---|---|---|---|
| MAS frequency/ kHz | 17 | 17 | 20 | 17 | 20 |
| Field/ T | 11.7 | 20.0 | 28.2 | 20.0 | 28.2 |
| Transfer I | HC-CP | HC-CP | HC-CP | HC-CP | HC-CP |
| $^{1}H$ field/ kHz | 60 | 60 | 55 | 60 | 75 |
| X field/ kHz | 43 | 43 | 29 | 43 | 48 |
| Shape | Tangent $^{1}H$ | Tangent $^{1}H$ | Tangent $^{1}H$ | Tangent $^{1}H$ | Tangent $^{1}H$ |
| $^{13}C$ carrier/ ppm | 100 | 100 | 100 | 100 | 100 |
| Time/ ms | 0.5 | 0.5 | 0.6 | 0.6 | 0.6 |
| Transfer II | DARR | DARR | DARR | DARR | DARR |
| $^{1}H$ field/ kHz | 17 | 17 | 20 | 17 | 20 |
| Carrier/ ppm | 100 | 100 | 100 | 100 | 100 |
| Time/ ms | 20 | 20 | 20 | 20 | 20 |
| t1 increments | 2560 | 2560 | 2560 | 2560 | 2560 |
| Sweep width (t1)/ kHz | 100 | 100 | 100 | 100 | 100 |
| Acquisition time (t1)/ ms | 12.8 | 12.8 | 12.8 | 12.8 | 12.8 |
| t2 increments | 3072 | 3072 | 3072 | 3072 | 3072 |
| Sweep width (t2)/ kHz | 100 | 100 | 100 | 100 | 100 |
| Acquisition time (t2)/ ms | 15.4 | 15.4 | 15.4 | 15.4 | 15.4 |
| $^{1}H$ Spinal64 decoupling power/ kHz | 90 | 90 | 90 | 90 | 90 |
| Interscan delay/ s | 2.7 | 2.7 | 2.7 | 2.7 | 2.7 |
| Number of scans | 12 | 12 | 12 | 4 | 4 |
| Measurement time/ h | 23 | 23 | 23 | 8 | 8 |

**Continue of table 1**: NMR parameters of 2D $^{13}C$-$^{13}C$ DARR correlation spectra recorded at 850 MHz (20.0 T) and 1200 MHz (28.2 T).

| **Experiment** | **DARR (TmcA)** | | **DARR (Pili)** | | **DARR (Nakedna virus)** | |
|---|---|---|---|---|---|---|
| MAS frequency/ kHz | 17 | 20 | 17 | 20 | 17 | 20 |
| Field/ T | 20.0 | 28.2 | 20.0 | 28.2 | 20.0 | 28.2 |
| Transfer I | HC-CP | HC-CP | HC-CP | HC-CP | HC-CP | HC-CP |
| $^{1}H$ field/ kHz | 60 | 70 | 60 | 70 | 60 | 75 |
| X field/ kHz | 43 | 44 | 43 | 44 | 41 | 47 |
| Shape | Tangent $^{1}H$ | Tangent $^{1}H$ | Tangent $^{1}H$ | Tangent $^{1}H$ | Tangent $^{1}H$ | Tangent $^{1}H$ |
| $^{13}C$ carrier/ ppm | 100 | 100 | 100 | 100.5 | 100 | 100 |
| Time/ ms | 0.5 | 0.5 | 0.7 | 1.0 | 0.6 | 0.6 |
| Transfer II | DARR | DARR | DARR | DARR | DARR | DARR |
| $^{1}H$ field/ kHz | 17 | 20 | 17 | 20 | 17 | 20 |
| Carrier/ ppm | 100 | 100 | 100 | 100 | 100 | 100 |
| Time/ ms | 20 | 20 | 20 | 20 | 20 | 20 |
| t1 increments | 2560 | 2560 | 2560 | 2560 | 2560 | 2560 |
| Sweep width (t1)/ kHz | 100 | 100 | 100 | 100 | 100 | 100 |
| Acquisition time (t1)/ ms | 12.8 | 12.8 | 12.8 | 12.8 | 12.8 | 12.8 |
| t2 increments | 3072 | 3072 | 3072 | 3072 | 3072 | 3072 |
| Sweep width (t2)/ kHz | 100 | 100 | 100 | 100 | 100 | 100 |
| Acquisition time (t2)/ ms | 15.4 | 15.4 | 15.4 | 15.4 | 15.4 | 15.4 |
| $^{1}H$ Spinal64 decoupling power/ kHz | 90 | 90 | 90 | 90 | 90 | 90 |
| Interscan delay/ s | 2.7 | 2.7 | 2.7 | 2.7 | 2.7 | 2.7 |
| Number of scans | 12 | 12 | 8 | 8 | 8 | 8 |
| Measurement time/ h | 23 | 23 | 16 | 16 | 16 | 16 |

**Supplementary table 2**: NMR parameters of 2D hNH and hCH spectra recorded on $^{13}C$-$^{15}N$ Rpo4/7 protein complex sample at 850 MHz (20.0 T) and 1200 MHz (28.2 T).

| **Protein** | **Rpo4/7 (hNH)** | | **Rpo4/7 (hCH)** | |
|---|---|---|---|---|
| Field / T | 20 | 28 | 20 | 28 |
| MAS frequency / kHz | 100 | 100 | 100 | 100 |
| Number of scans | 48 | 48 | 40 | 40 |
| t1 increment | 524 | 740 | 724 | 1024 |
| Sweep width (t1) / ppm | 120 | 120 | 80 | 80 |
| Acquisition time (t1) / ms | 25.3 | 25.3 | 21.2 | 21.2 |
| t2 increment | 4096 | 4096 | 4096 | 4096 |
| Sweep width (t2) / ppm | 46.7 | 46.2 | 46.7 | 46.2 |
| Acquisition time (t2) / ms | 51.6 | 36.8 | 25.8 | 36.8 |
| $^{1}H$ dec (swftPPM) / kHz | 10 | 10 | 10 | 10 |
| $^{15}N$ dec (WALTZ64) / kHz | 5 | 5 | - | - |
| $^{13}C$ dec (WALTZ64) / kHz | - | - | 5 | 5 |
| Water sup. (120 ms) / kHz | 20 | 20 | 20 | 20 |
| Interscan delay / s | 1.27 | 1.65 | 1.27 | 1.65 |
| Experiment time | 10h10 | 18h | 11h30 | 20h50 |
| **Transfer 1** | HN (dipolar) | HN (dipolar) | HC (dipolar) | HC (dipolar) |
| $^{1}H$ field / kHz | 83.8 | 85 | 83.8 | 76.6 |
| $^{15}N$ or $^{13}C$ field / kHz | 13.9 | 22 | 18.5 | 20 |
| Shape | Tangent $^{1}H$ | Tangent $^{1}H$ | Tangent $^{1}H$ | Tangent $^{1}H$ |
| Time / ms | 1.4 | 1.4 | 0.7 | 0.7 |
| **Transfer 2** | NH (dipolar) | NH (dipolar) | CH (dipolar) | CH (dipolar) |
| $^{1}H$ field / kHz | 80.6 | 86.6 | 73.5 | 76.6 |
| $^{15}N$ or $^{13}C$ field / kHz | 13.9 | 22 | 20 | 20 |
| Shape | Tangent $^{1}H$ | Tangent $^{1}H$ | Tangent $^{1}H$ | Tangent $^{1}H$ |
| Time / ms | 1.0 | 0.9 | 0.7 | 0.7 |
| Carrier $^{15}N$ / ppm | 117.5 | 177.5 | - | - |
| Carrier $^{13}C$ / ppm | - | - | 40 | 40 |
| Carrier $^{1}H$ / ppm | 4.8 | 4.8 | 4.8 | 4.8 |

**Supplementary table 3**: NMR parameters of 2D hNH and hCH spectra recorded on $^{13}C$-$^{15}N$ ASC sample at 850 MHz (20.0 T) and 1200 MHz (28.2 T).

| **Protein** | **ASC (hNH)** | | **ASC (hCH)** | |
|---|---|---|---|---|
| Field / T | 20 | 28 | 20 | 28 |
| MAS frequency / kHz | 100 | 100 | 100 | 100 |
| Number of scans | 64 | 64 | 32 | 32 |
| t1 increment | 512 | 724 | 1536 | 2170 |
| Sweep width (t1) / ppm | 70 | 70 | 180 | 180 |
| Acquisition time (t1) / ms | 42.5 | 42.5 | 20 | 20 |
| t2 increment | 2048 | 3072 | 2048 | 3072 |
| Sweep width (t2) / ppm | 20 | 19.8 | 46.7 | 46.3 |
| Acquisition time (t2) / ms | 60.2 | 64.5 | 25.8 | 27.6 |
| $^{1}H$ dec (swftPPM) / kHz | 10 | 10 | 10 | 10 |
| $^{15}N$ dec (WALTZ64) / kHz | 5 | 5 | - | - |
| $^{13}C$ dec (WALTZ64) / kHz | - | - | 5 | 5 |
| Water sup. (120 ms) / kHz | 20 | 20 | 20 | 20 |
| Interscan delay / s | 1 | 1.18 | 1 | 1.18 |
| Experiment time | 11h | 18h41 | 15h51 | 27h03 |
| **Transfer 1** | HN (dipolar) | HN (dipolar) | HC (dipolar) | HC (dipolar) |
| $^{1}H$ field / kHz | 67.4 | 89.4 | 65.8 | 132.7 |
| $^{15}N$ or $^{13}C$ field / kHz | 20.0 | 14.7 | 30.4 | 32.9 |
| Shape | Tangent $^{1}H$ | Tangent $^{1}H$ | Tangent $^{1}H$ | Tangent $^{1}H$ |
| Time / ms | 1.5 | 1.3 | 0.7 | 0.7 |
| **Transfer 2** | NH (dipolar) | NH (dipolar) | CH (dipolar) | CH (dipolar) |
| $^{1}H$ field / kHz | 65.8 | 88.0 | 58.8 | 130.9 |
| $^{15}N$ or $^{13}C$ field / kHz | 20.0 | 14.7 | 30.4 | 32.9 |
| Shape | Tangent $^{1}H$ | Tangent $^{1}H$ | Tangent $^{1}H$ | Tangent $^{1}H$ |
| Time / ms | 1.5 | 1.3 | 0.7 | 0.7 |
| Carrier $^{15}N$ / ppm | 107 | 106.4 | - | - |
| Carrier $^{13}C$ / ppm | - | - | 58.5 | 58.0 |
| Carrier $^{1}H$ / ppm | 4.8 | 4.8 | 4.4 | 4.4 |

**Supplementary table 4**: NMR parameters of 2D hNH spectra recorded on the deuterated samples (dCp149 and dNS4B) at 850 MHz (20.0 T) and 1200 MHz (28.2 T).

| **Protein** | **dCp149** | | **dNS4B** | |
|---|---|---|---|---|
| Field / T | 20 | 28 | 20 | 28 |
| MAS frequency / kHz | 100 | 100 | 100 | 100 |
| Number of scans | 48 | 48 | 80 | 80 |
| t1 increment | 362 | 512 | 284 | 400 |
| Sweep width (t1) / ppm | 80 | 80 | 40 | 40 |
| Acquisition time (t1) / ms | 26.2 | 26.2 | 41.2 | 41.1 |
| t2 increment | 4096 | 4096 | 4096 | 4096 |
| Sweep width (t2) / ppm | 46.7 | 46.2 | 46.7 | 46.2 |
| Acquisition time (t2) / ms | 51.6 | 36.8 | 51.6 | 36.8 |
| $^{1}$H dec (swftPPM) / kHz | 10 | 10 | 10 | 10 |
| $^{15}$N dec (WALTZ64) / kHz | 5 | 5 | 5 | 5 |
| Water sup. (120 ms) / kHz | 20 | 20 | 20 | 20 |
| Interscan delay / s | 1.5 | 2.2 | 1.2 | 1.43 |
| Experiment time | 8h | 16h30 | 8h50 | 14h22 |
| **Transfer 1** | HN (dipolar) | HN (dipolar) | HN (dipolar) | HN (dipolar) |
| $^{1}$H field / kHz | 79 | 81 | 82.2 | 87.5 |
| $^{15}$N field / kHz | 14 | 14 | 14 | 14.6 |
| Shape | Tangent $^{1}$H | Tangent $^{1}$H | Tangent $^{1}$H | Tangent $^{1}$H |
| Time / ms | 1.6 | 1.5 | 1.0 | 1.0 |
| **Transfer 2** | NH (dipolar) | NH (dipolar) | NH (dipolar) | NH (dipolar) |
| $^{1}$H field / kHz | 75.7 | 78.1 | 78.7 | 87.5 |
| $^{15}$N field / kHz | 14 | 14 | 14 | 14.6 |
| Shape | Tangent $^{1}$H | Tangent $^{1}$H | Tangent $^{1}$H | Tangent $^{1}$H |
| Time / ms | 1.8 | 1.7 | 1.2 | 1.2 |
| Carrier $^{15}$N / ppm | 107 | 107 | 116.5 | 116.5 |
| Carrier $^{1}$H / ppm | 4.8 | 4.8 | 4.8 | 4.8 |

**Supplementary table 5**: amide proton transverse relaxation times $T_2'$ ($^1H_N$) measured in $^1$H-detected experiments at 850 MHz (20.0 T) and 1200 MHz (28.2 T).

| **Protein** | **dCp149** | | **dNS4B** | | **Rpo4/7** | | **ASC** | |
|---|---|---|---|---|---|---|---|---|
| Field / T | 20.0 | 28.2 | 20.0 | 28.2 | 20.0 | 28.2 | 20.0 | 28.2 |
| $T_2'$ ($^1H_N$) / ms | 10.40 | 11.36 | 4.69 | 4.73 | 2.53 | 2.69 | 2.6 | 2.77 |

### Supplementary References:
